# Chromosomal organization of biosynthetic gene clusters suggests plasticity of myxobacterial specialized metabolism including descriptions for nine novel species: *Archangium lansinium* sp. nov., *Myxococcus landrumus* sp. nov., *Nannocystis bainbridgea* sp. nov., *Nannocystis poenicansa* sp. nov., *Nannocystis radixulma* sp. nov., *Polyangium mundeleinium* sp. nov., *Pyxidicoccus parkwaysis* sp. nov., *Sorangium aterium* sp. nov., *Stigmatella ashevillena* sp. nov

**DOI:** 10.1101/2023.03.08.531766

**Authors:** Andrew Ahearne, Kayleigh Phillips, Thomas Knehans, Miranda Hoing, Scot E. Dowd, D. Cole Stevens

## Abstract

Natural products discovered from bacteria provide critically needed therapeutic leads for drug discovery, and myxobacteria are an established source for metabolites with unique chemical scaffolds and biological activities. Myxobacterial genomes accommodate an exceptional number and variety of biosynthetic gene clusters (BGCs) which encode for features involved in specialized metabolism. Continued discovery and sequencing of novel myxobacteria from the environment provides BGCs for the genome mining pipeline. Herein, we describe the collection, sequencing, and genome mining of 20 myxobacteria isolated from rhizospheric soil samples collected in North America. Nine isolates where determined to be novel species of myxobacteria including representatives from the genera *Archangium, Myxococcus, Nannocystis, Polyangium, Pyxidicoccus, Sorangium*, and *Stigmatella*. Growth profiles, biochemical assays, and descriptions are provided for all proposed novel species. We assess the BGC content of all isolates and observe differences between Myxococcia and Polyangiia clusters. Utilizing complete or near complete genome sequences we compare the chromosomal organization of BGCs of related myxobacteria from various genera and suggest spatial proximity of hybrid, modular clusters contributes to the metabolic adaptability of myxobacteria.

## INTRODUCTION

Myxobacteria are metabolically “gifted” bacteria with large genomes accommodating an exceptional number of biosynthetic gene clusters (BGCs) and the potential to produce highly diverse specialized metabolites (1–4). Excellent reservoirs of candidate therapeutics, over 100 unique metabolite scaffolds have been discovered from myxobacteria (2). Extensive metabolomic analysis of ~2,300 myxobacterial extracts revealed a correlation between detected metabolites and taxonomic distance with genus-level hierarchical clustering of metabolite profiles (5). Although limited to metabolic profiles of axenically grown myxobacteria, this observation suggests investigation of lesser-studied genera within the phylum Myxococcota might increase the likelihood of metabolite discovery. Ongoing natural product discoveries from novel species of myxobacteria reinforce the need for continued isolation and characterization of environmental myxobacteria (6–9). Most recently described Myxococcota belong to the genera *Corallococcus, Myxococcus*, and *Pyxidicoccus* (10–14), and comparatively fewer members of lesser-studied myxobacterial taxa have been reported over the last decade (15–17). For example, no new type strain *Stigmatella* has been reported since 2007. Herein we report isolation and genome sequencing of 20 environmental myxobacteria including representatives from the less-well-studied *Archangium, Nannocystis*, and *Polyangium*. Complete and near-complete genome data enabled a thorough assessment of BGC content which revealed 1) significant differences in cluster sizes of Myxococcia and Polyangiia, 2) unique biosynthetic capacity of *Nannocystis*, and 3) chromosomal organization of myxobacterial BGCs.

## RESULTS

### Isolation and genomic comparison of 20 isolated myxobacteria

Rhizospheric soil samples collected from shrubs and trees were screened for bacterial swarms using standard prey-baiting and filter paper degradation methods (18–20) to isolate environmental myxobacteria (Supplemental Figure 1)(21). Morphology screening of visible swarms facilitated isolation of myxobacteria from multiple genera with a specific focus on lesser-studied myxobacteria. A total of 20 environmental isolates of putative myxobacteria including eight agarolytic isolates were obtained as monocultures (Figure 1). Lesser-studied myxobacteria include genera with agarolytic phenotypes such as *Nannocystis, Polyangium*, and *Sorangium* thus numerous agarolytic isolates with similar morphologies were advanced for genome sequencing (18). We have previously discussed four of the 20 environmental isolates (SCHIC03, SCPEA02, NCCRE02, NCSPR01) (22). Genome sequencing of all isolates provided five complete genomes, seven draft genomes with ≤3 contigs, three draft genomes with 5-8 contigs, and five lower quality genome assemblies with ≤44 contigs (Table 1). Genome sizes ranged from 9,459,689-13,831,693 Mb and GC content varied from 68.1-71.5%. High quality assemblies enabled subsequent whole genome comparison approaches for phylogenetic analysis and assessment of biosynthetic gene cluster content and organization.

**Figure 1:**
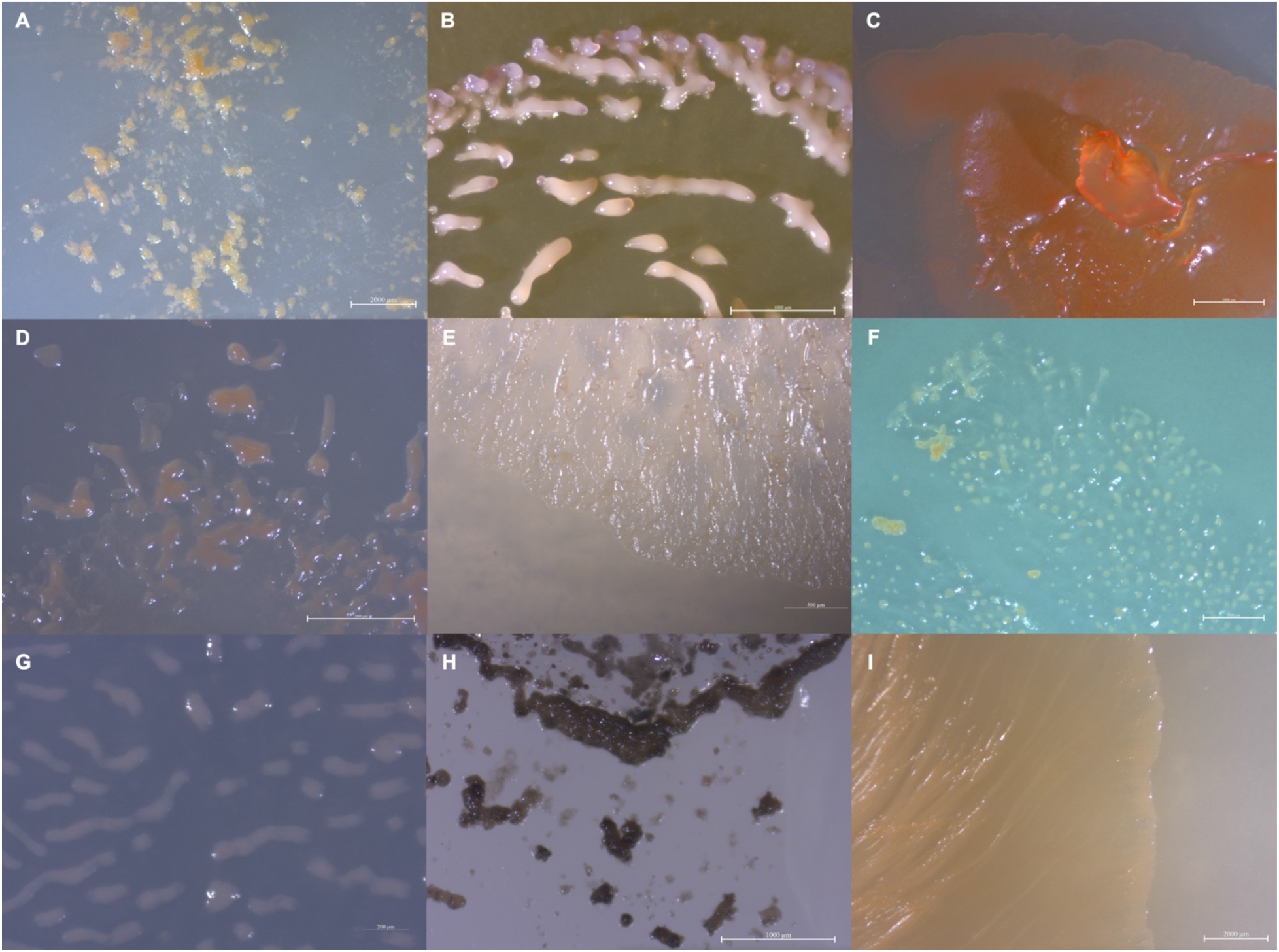
Images depicting the observed variety in isolate morphology A) MIWBW, B) SCHIC03, C) BB15-2, D) FL3, E) NCELM, F) RJM3, G) SCPEA02, H) WIWO2, I) NCWAL01.

**Table 1:** Genome assembly data for sequenced isolates with proposed novel species bolded. Asterisks denote previously reported isolates.

| <u>Isolate</u> | <u>Size (bp)</u> | <u>CDS</u> | <u>GC%</u> | <u>N50</u> | <u>L50</u> | <u>Contigs</u> |
| --- | --- | --- | --- | --- | --- | --- |
| **MIWBW** | 13,831,693 | 11,592 | 68.1 | 13,758,192 | 1 | 2 |
| **SCHIC03\*** | 10,367,529 | 8,339 | 68.6 | - | 1 | 1 |
| **BB15-2** | 11,048,555 | 9,803 | 71.5 | 3,033,751 | 2 | 5 |
| **FL3** | 13,228,663 | 12,183 | 71.5 | - | 1 | 1 |
| **NCELM** | 12,940,226 | 11,412 | 71.2 | 791,632 | 7 | 29 |
| **RBIL** | 12,529,437 | 11,262 | 71.4 | 6,343,246 | 1 | 3 |
| **RJM3** | 13,254,631 | 11,155 | 68.6 | 13,235,919 | 1 | 2 |
| **SCPEA02\*** | 13,211,253 | 10,588 | 69.6 | - | 1 | 1 |
| **WIWO2** | 13,472,481 | 11,616 | 71.1 | 2,525,385 | 2 | 7 |
| **NCWAL01** | 10,951,030 | 8,884 | 66.9 | 941,459 | 1 | 2 |
| NCSPR01* | 9,785,177 | 8,033 | 70.1 | 9,343,940 | 1 | 3 |
| NCCRE02* | 10,538,407 | 8,589 | 69.7 | 3,024,381 | 2 | 8 |
| MSG2 | 13,340,839 | 10,353 | 70.3 | 13,193,640 | 1 | 2 |
| NCRR | 9,787,125 | 8,044 | 70.2 | - | 1 | 1 |
| MISCRS | 10,873,373 | 8,896 | 70.2 | 10,790,239 | 1 | 2 |
| SCPEA4 | 12,461,442 | 11,002 | 71.3 | 1,797,159 | 3 | 22 |
| BB11-1 | 9,459,689 | 7,665 | 70.6 | 427,033 | 7 | 44 |
| BB12-1 | 10,337,840 | 8,427 | 69.6 | 593,731 | 6 | 38 |
| ILAH1 | 12,844,680 | 11,575 | 71.2 | 573,330 | 8 | 41 |
| NMCA1 | 9,502,182 | 8,152 | 69 | - | 1 | 1 |

### Phylogenetic relationships of isolated myxobacteria

Initial phylogenetic analysis using 16S rRNA sequences of type strain myxobacteria (obtained from the List of Prokaryotic names with Standing in Nomenclature (LPSN)) suggested the environmental isolates included 1 *Archangium*, 5 *Corallococcus*, 3 *Myxococcus*, 6 *Nannocystis*, 1 *Polyangium*, 2 *Pyxidicoccus*, 1 *Sorangium*, and 1 *Stigmatella* (Supplemental Figure 2). Utilizing genome data from each isolate, conserved gene sequence similarities between isolates and type strain myxobacteria were determined using average nucleotide identity (ANI) and digital DNA-DNA hybridization values (dDDH) according to established methods for taxonomic assignment of myxobacteria (10, 11). Resulting ANI and dDDH values indicated ten of the 20 environmental isolates to be novel species with values below the respective cutoffs of 95% and 70% when compared to most similar type strains (Figure 2 and Supplemental Table 1). Isolate MIWBW is most phylogenetically similar to *Archangium gephyra* DSM2261^T^ and *A. gephyra* Cbvi76 (previously referred to as *Cystobacter violaceus* Cbvi76 (23)) (Figure 2A). Of the four published type strain *Archangium, A. gephyra* DSM2261^T^ is currently the only representative sufficiently sequenced for comparative genomic analysis. More rigorous analysis comparing sequenced representatives of closely related *Cystobacter* and *Melittangium* reinforced MIWBW as a novel species. This analysis also revealed *Cystobacter gracilis* DSM 14753^T^ to be an outlier within the three genera with ANI values below 77.5 for all included representatives. Isolate SCHIC03 is most phylogenetically similar to *Myxococcus stipitatus* DSM 14675^T^ when compared to eight *Myxococcus* type strains (Figure 2B). Also observed by Chambers *et al*., the ANI value between established type strain species *Myxococcus xanthus* DSM 16526^T^ and *Myxococcus virescens* DSM 2260^T^ is above the threshold for novel species (11). Initial 16S rRNA analysis suggested environmental isolates RBIL2, FL3, BB15-2, and NCELM to all be novel *Nannocystis* species. However, of the three type strain *Nannocystis*, there was no genome data for *Nannocystis pusilla* DSM 14622^T^ (also referred to as *N. pusilla* Na p29^T^). Subsequent sequencing of *N. pusilla* DSM 14622^T^ and comparison including our genome data for *N. pusilla* DSM 14622^T^ revealed RBIL2 to be a subspecies of *N. pusilla* that is slightly above the novel species threshold (Figure 2E). Isolates BB15-2, FL3, and NCELM are most phylogenetically similar to *Nannocystis exedans* DSM 71^T^ and are significantly distinct from each other. Our proposed addition of three *Nannocystis* doubles the current member total. Isolate RJM3 is most phylogenetically similar to *Polyangium fumosum* DSM 14688^T^ (Figure 2C). However, only three of ten *Polyangium* type strains have sufficient 16S rRNA and genome sequence data suitable for thorough analysis (15, 24). Isolate SCPEA02 is most phylogenetically similar to *Pyxidicoccus caerfyrddinensis* CA032A^T^ (Figure 2G) (11). The proposed addition to the *Pyxidicoccus* will make SCPEA02 just the fourth type strain *Pyxidicoccus*. Isolate WIWO2 is most phylogenetically similar to *Sorangium cellulosum* Soce56, but no type strain of *Sorangium* have been sufficiently sequenced for comparative genomics (Figure 2D). Alternatively, 16S RNA analysis suggests WIWO2 is most phylogenetically similar to *Sorangium kenyense* Soce 375^T^ (Supplemental Figure 2). Isolate NCWAL01 is most phylogenetically similar to *Stigmatella aurantiaca* DSM17044^T^ and *St. aurantiaca* DW4_3-1. Interestingly, our analysis indicates ANI values above the threshold for novel species for all three type strains of *Stigmatella* (Figure 2F).

**Figure 2:**
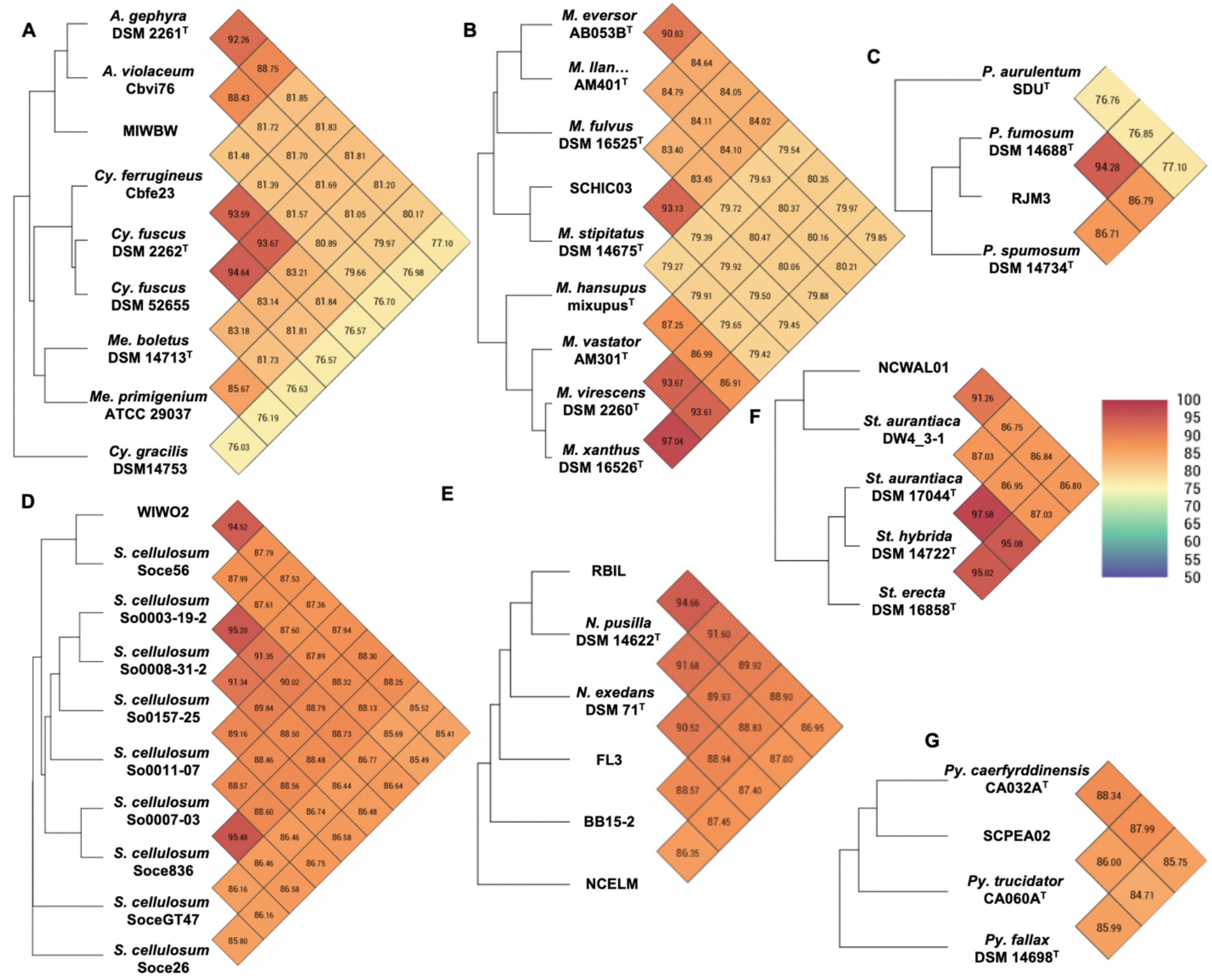
Heatmaps generated from OrthoANI values calculated using OAT for A) MIWBW and sequenced strains of *Archangium, Cystobacter*, and *Melitangium* B) SCHIC03 and type strain *Myxococcus*, C) RJM3 and type strain *Polyangium*, D) WIWO2 and sequenced *Sorangium*, E) RBIL2, FL3, BB15-2, NCELM, and sequenced *Nannocystis*, F) NCWAL01 and sequenced *Stigmatella*, and G) SCPEA02 and type strain *Pyxidicoccus*. The established cutoff for new species designation is <95%. *“M. llan...”* is an abbreviation of *Myxococcus* *llanfairpwllgwyngyllgogerychwyrndrobwllllantysiliogogogochensis* AM401^T^.

Environmental isolates NCSPR01 and NCRR are both highly similar subspecies of *Corallococcus coralloides* DSM 2259^T^ (Supplemental Table 1 and Supplemental Figure 3A). As previously suggested by Ahearne et al., isolate NCCRE02 is a subspecies of *Corallococcus exiguus* DSM 14696^T^ (Supplemental Table 1 and Supplemental Figure 3A) (22). Isolate BB12-1 is likely a subspecies of *Corallococcus terminator* CA054A^T^, and isolate BB11-1 is potentially a novel species of *Corallococcus* (Supplemental Table 1 and Supplemental Figure 3B). However, fragmented genome assemblies for BB11-1 and BB12-1 limited our confidence in precise taxonomic placement. Isolate MISCRS is a subspecies of *Myxococcus fulvus* DSM 16525^T^, and isolate NMCA is a subspecies of *M. xanthus* DSM 16526^T^ (Supplemental Table 1 and Supplemental Figure 3C). Isolate MSG2 is a subspecies of *Py. caerfyrddinensis* CA032A^T^ (Supplemental Table 1 and Supplemental Figure 3D). Isolate SCPEA04 is a *Nannocystis* highly similar to NCELM, and isolates RBIL2 and ILAH1 are both subspecies of *N. pusilla* DSM 14622^T^ (Supplemental Table 1 and Supplemental Figure 3E).

### Physiological and biochemical analysis of nine novel genomospecies

All isolated strains swarmed on VY/2 media, and growth characteristics at various pH values and temperatures were analyzed for all nine novel species (Table 2). All nine strains grew at 25-30°C, and SCPEA02 grew at temperatures up to 40°C. Growth at pH 7 was observed from all strains, and SCHIC03, NCELM, and SCPEA02 all grew at pH 6-9. Agarolytic strains include BB15-2, WIWO2, FL3, NCELM, and RJM3. Metabolic activity was assessed for all strains (Table 3), and none were able to reduce nitrate or metabolize arginine, glucose, or urea. All strains were able to hydrolyze esculin, and all except FL3 and WIWO2 hydrolyzed gelatin. SCPEA02 and SCHIC03 were the only strains that did not exhibit alkaline phosphatase activity. MIWBW was the only strain to demonstrate both trypsin and α-chymotrypsin activity and possessed overlapping characteristics with *A. gephyra* (25). Growth and activity of SCHIC03 was most similar to *M. stipitatus* and *M. fulvus* (11). Growth profiles and biochemical activities of BB15-2, FL3, and NCELM are similar to other *Nannocystis;* however, all three demonstrated comparatively limited pH-dependent growth (17). Unlike *Nannocystis konarekensis, N. exedans*, and *N. pusilla*, none of the *Nannocystis* strains grew at pH 10. Temperature and pH-dependent growth ranges for RJM3 are notably different from other *Polyangium*, which all grow at temperatures above 30°C and pH ranges of 6-8.5 (15). The growth profile and biochemical activity of SCPEA02 were closely aligned with the reported activities of *Py. caerfyrddinensis* (11). Growth profile and biochemical activity of WIWO2 overlapped somewhat with the recently described *Sorangium* species (16). Characterization and description of all type strain *Stigmatella* pre-date modern description methodology, however *St. aurantiaca* and NCELM have similar growth and biochemical profiles (26, 27).

**Table 2:** Growth characteristics for isolates proposed to be novel species. Symbols indicate growth using percent increase in swarm diameters over ten days with “-” as no growth, “+” low growth (≤39% swarm diameter), “++” moderate growth (40-75% swarm diameter), and “+++” high growth (76-100% swarm diameter).

| temperature<br>(°C) | WIWO2 | BB15-2 | FL3 | RJM3 | NCWAL01 | SCHIC03 | NCELM | SCPEA02 | MIWBW |
| --- | --- | --- | --- | --- | --- | --- | --- | --- | --- |
| 20 | - | ++ | + | + | +++ | ++ | - | + | ++ |
| 25 | + | +++ | +++ | +++ | +++ | +++ | ++ | +++ | +++ |
| 30 | +++ | ++ | +++ | +++ | +++ | ++ | +++ | +++ | +++ |
| 35 | + | - | +++ | - | + | +++ | - | +++ | +++ |
| 40 | - | - | - | - | - | - | - | +++ | - |
| pH |  |  |  |  |  |  |  |  |  |
| 5 | - | - | - | - | - | - | - | - | - |
| 6 | + | - | + | - | ++ | +++ | ++ | ++ | + |
| 7 | +++ | +++ | +++ | +++ | +++ | +++ | +++ | +++ | +++ |
| 8 | - | +++ | +++ | - | +++ | ++ | +++ | + | +++ |
| 9 | - | - | - | - | - | + | + | + | - |
| 10 | - | - | - | - | - | ++ | - | - | - |

**Table 3:**
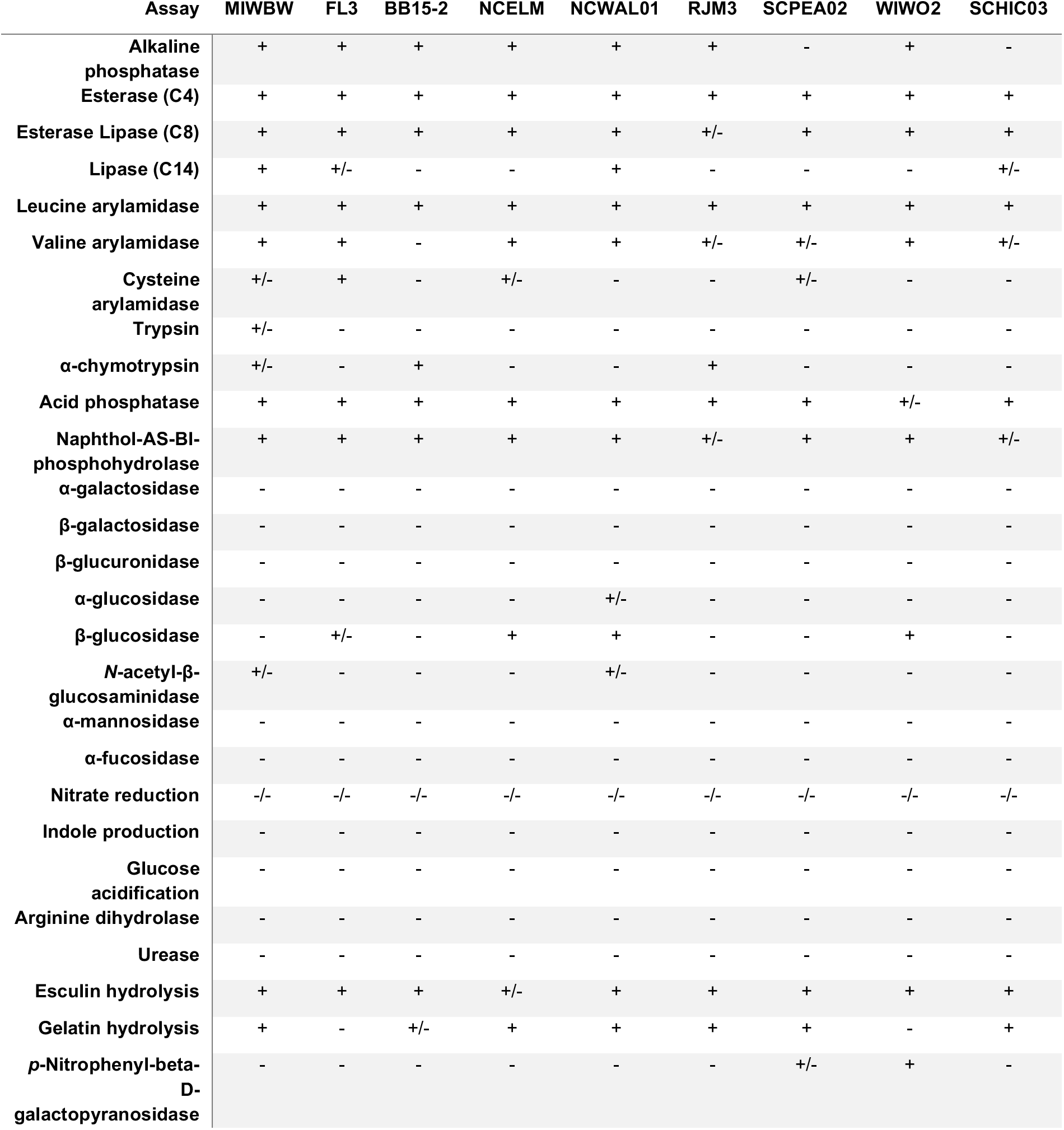
Enzymatic activity data for isolates proposed to be novel species. Symbols indicate “-” no activity, “+/-” low activity, and “+” high activity.

### Biosynthetic potential of myxobacterial isolates

AntiSMASH analysis of BGCs from all sequenced isolates provided notable differences in BGC contents, sizes, and similarities with previously characterized clusters (28, 29). A total of 735 BGCs were predicted from 20 genome assemblies and only 36 were identified by antiSMASH to be fragmented clusters (~5%) with the vast majority of fragmented BGCs included in sequenced *Corallococcus* strains (20 fragmented BGCs). Genome data from sequenced *Archangium*, *Myxococcus*, *Polyangium*, *Pyxidicoccus*, *Sorangium*, and *Stigmatella* provided three or fewer fragmented BGCs total from each genus. Of our isolated strains MIWBW had the most predicted BGCs with 52 total (zero fragmented), and NMCA1 had the least with 23 total (zero fragmented). The average length of BGCs from all 20 sequenced isolates was ~56 kb. Clusters from *Nannocystis* strains were significantly shorter (average size ~30 kb) than *Corallococcus, Myxococcus*, and *Pyxidicoccus* clusters (Figure 3A). Clusters from *Pyxidicoccus* strains were significantly longer than *Archangium, Nannocystis, Polyangium*, and *Sorangium* clusters. Interestingly, Myxococcia clusters were significantly longer than Polyangiia BGCs (Figure 3B).

**Figure 3:**
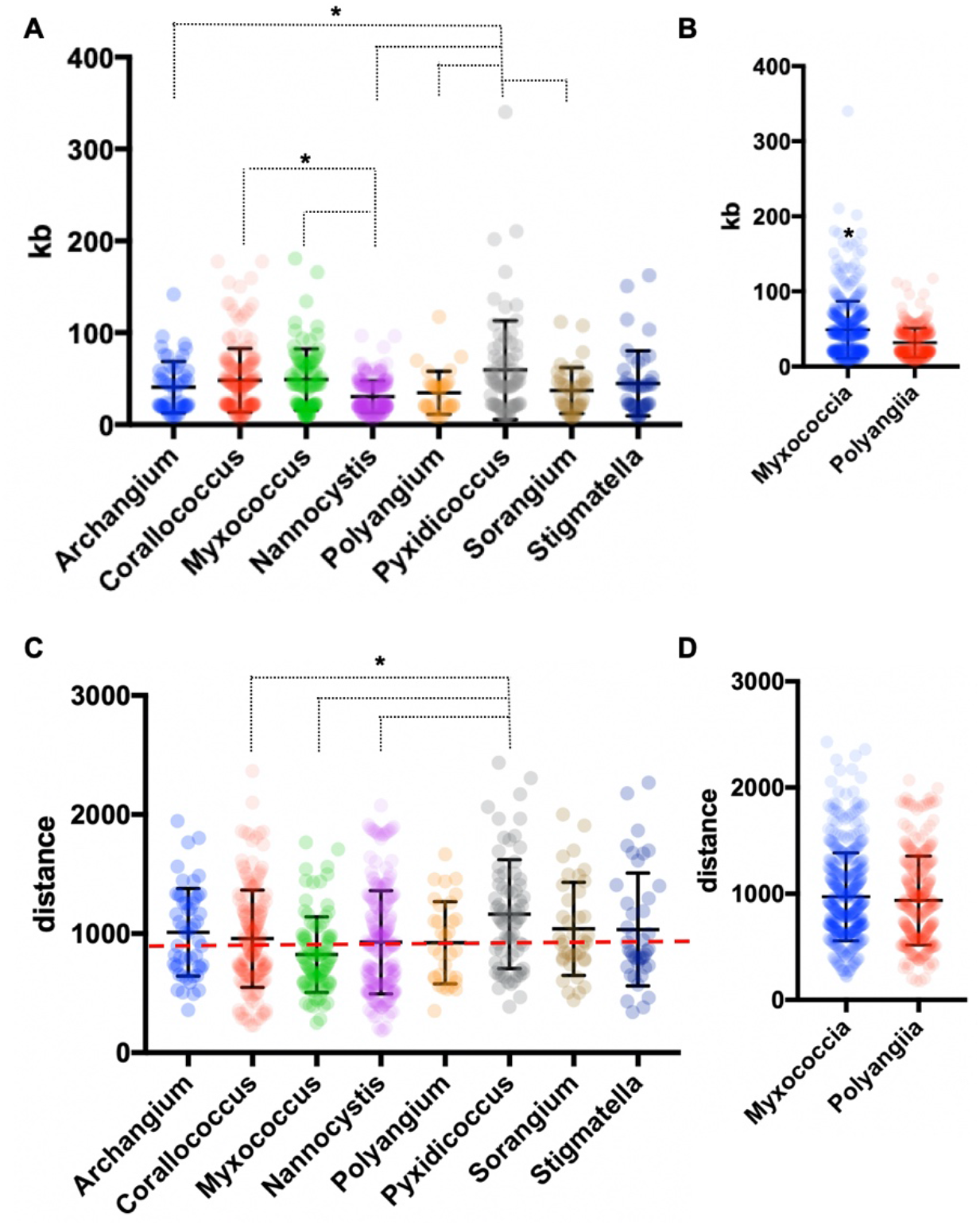
Genus-level analysis of BGCs and BiG-FAM distances. A) Genus- and (B) class-level comparison of BGC size. C) Genus- and (D) class-level comparison of BiG-FAM distances of BGCs with the red, dashed line indicating *d*=900. Ordinary one-way ANOVA with multiple comparisons used to determine significance for genus-level analysis, and Welch’s t test used to determine significance of class-level analysis (for genus-level analysis *Archangium* n=52, *Corallococcus* n=176, *Myxococcus* n=91, *Nannocystis* n=212, *Polyangium* n=32, *Pyxidicoccus* n=73, *Sorangium* n=39, *Stigmatella* n=42; *p*<0.05 and class-level analysis Myxococcia n=434, Polyangiia n=283; *p*<0.0001).

All identified BGCs were compared with the 1,225,071 BGCs and 29,955 gene cluster families (GCFs) included in the BiGFAM database (30, 31). Utilizing a previously established clustering threshold (T=900) to determine distance from database GCFs, we evaluated our 764 BGCs for similarity with BiGFAM BGCs. Clusters below the arbitrary threshold have similarities with BiGFAM GCFs and are likely less novel than clusters above the threshold (31, 32). Clusters from *Pyxidicoccus* strains (SCPEA02 and MSG2) had the highest average distance (1165) and somewhat predictably *Myxococcus* strains had the lowest (824) with an average distance below the threshold (Figure 3C). Average distance of *Pyxidicoccus* BGCs was significantly higher than the average distances of *Corallococcus, Myxococcus*, and *Nannocystis* clusters. Although *Nannocystis* and *Polyangium* are lesser-studied myxobacteria, average distances of clusters from members of each genus were just above the threshold for novelty (923 and 924 respectively). No significant difference was observed between the average distances of Myxococcia and Polyangiia clusters (Figure 3D).

Of the 764 predicted BGCs, 384 clusters (~50%) had distances above the threshold. Removal of clusters below the threshold revealed differences in remaining cluster types across genera (Figure 4). Subsequent comparison of these 384 BGCs with cluster similarities identified during antiSMASH analysis revealed 52 clusters were either highly homologous to characterized clusters deposited in the MiBIG database (33) or included embedded clusters with high similarity to known clusters. For example, the myxochelin BGC was found to be embedded in 10 clusters that scored above the BiGFAM threshold (34, 35). Although co-clustering likely impedes analysis of novelty and similarity to BiGFAM GCFs, we suggest such co-clustering does not necessarily preclude uniqueness of proximal clusters. This analysis also identified clusters that likely produce known metabolites including: 2-methylisoborneol (FL3), alkylpyrone 407/393 (BB11-1, BB12-1), aurafuron A (NCWAL01), carotenoid (RJM3), chloromyxamide (MSG2), dawenol (BB12-1, SCHIC03), dkxanthene (MISCRS, NMCA1, SCHIC03), geosmin (BB12-1, NCRR, NCSPR01), myxoprincomide (MSG2), nannocystin A (FL3), phenalamide A2 (SCHIC03), pyrronazol B (RBIL2), rhizopodin (NCWAL01, SCHIC03), ripostatin A/B/C (WIWO2), and VEPE/AEPE/TG-1 (BB11-1, BB12-1, CRE02, MIWBW, MSG2, NCRR) clusters (36–51). Modular clusters with high homology but differing organization that likely produce analogs of known metabolites were also identified including the 2-methylisoborneol (NCELM), fulvuthiacene A/B (MISCRS), lyngbyatoxin A (NCELM, SCPEA04), myxoprincomide (MISCRS, SCPEA02, SCHIC03, MIWBW), pyrronazol B (BB15-2, ILAH1) and violacein (SCHIC03) clusters (38, 46, 49, 52–55). Additional cluster similarity was identified across the eight sequenced *Nannocystis* strains. For example, strain SCPEA04 contained no unique clusters that were not also present in the other seven *Nannocystis* genomes, and no strain had more than five unique clusters. Five intriguing novel phosphonate clusters from four *Nannocystis* strains (BB15-2, NCELM, SCPEA04, RBIL2) and WIWO2 highlight the potential for novel metabolite discovery from the lesser-studied myxobacteria. Typically discovered from *Streptomyces* and *Pseudomonas* spp. (56, 57), no phosphonate metabolites have previously been discovered from a myxobacterium. Further differences between Myxococcia and Polyangiia cluster content was revealed with subsequent analysis of BGC relatedness between isolates and sequenced myxobacteria deposited in the antiSMASH database using BiG-SCAPE (Supplemental Figures 4 and 5). Resulting gene cluster families included connectivities between clusters of various Myxococcia suggesting an inherited overlap in specialized metabolism. However all *Nannocystis* and *Sorangium* gene cluster families with 2 or more clusters were exclusively genus-specific, and all BGCs from *Polyangium* sp. strain RJM3 were present in singleton gene cluster families.

**Figure 4:**
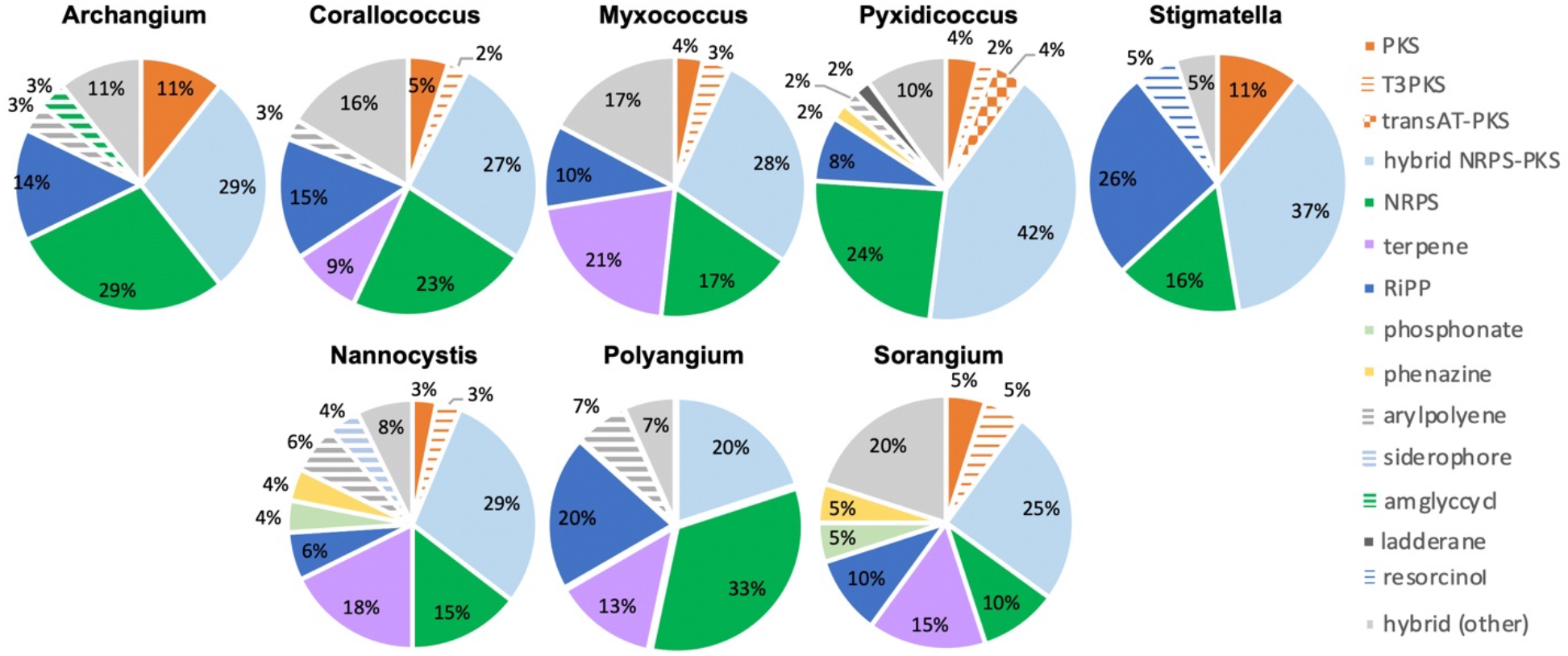
Genus-level distribution of likely novel BGCs by cluster type with cluster novelty assumed for BGCs with distances below the BiG-FAM threshold (*d*=900). Cluster type designations provided by antiSMASH analysis.

Additional analysis of myxobacterial BGCs that encode metabolites with reported ecological utility unveiled notable differences and similarities amongst genera. The myxochelin cluster is present in all sequenced strains excluding WIWO2 and *Nannocystis* strains (Figure 5). Myxochelin has been discovered from numerous myxobacteria and functions as a siderophore during iron starvation conditions (58). One or more alternative siderophore clusters are present in all *Nannocystis* strains. Carotenoid and VEPE/AEPE/TG-1 clusters were present in all analyzed Myxococcia and notably absent in all Polyangiia. Geosmin serves as a small molecule deterrent or “warning signal” to dissuade predatory nematodes, and the geosmin cluster is present in all strains (59). Myxovirescin (60, 61) and myxoprincomide (62) benefit *M. xanthus* predation of Gram negative and Gram positive prey respectively (63). Myxovirescin has been found to significantly benefit *M. xanthus* predation of *E. coli* (61,64). However, none of the investigated strains including the *M. xanthus* strain NMCA1 possessed a cluster with similarity to the myxovirescin BGC. Aside from the absence of a myxovirescin cluster, NMCA1 and *M. xanthus* DK1622 share incredible similarity in BGC content including high similarity surrounding the myxovirescin BGC in *M. xanthus* DK1622 (Supplemental Figure 6). The myxoprincomide cluster or a cluster with high similarity to it is present in all strains excluding members of the class Polyangiia and NCWAL01.

**Figure 5:**
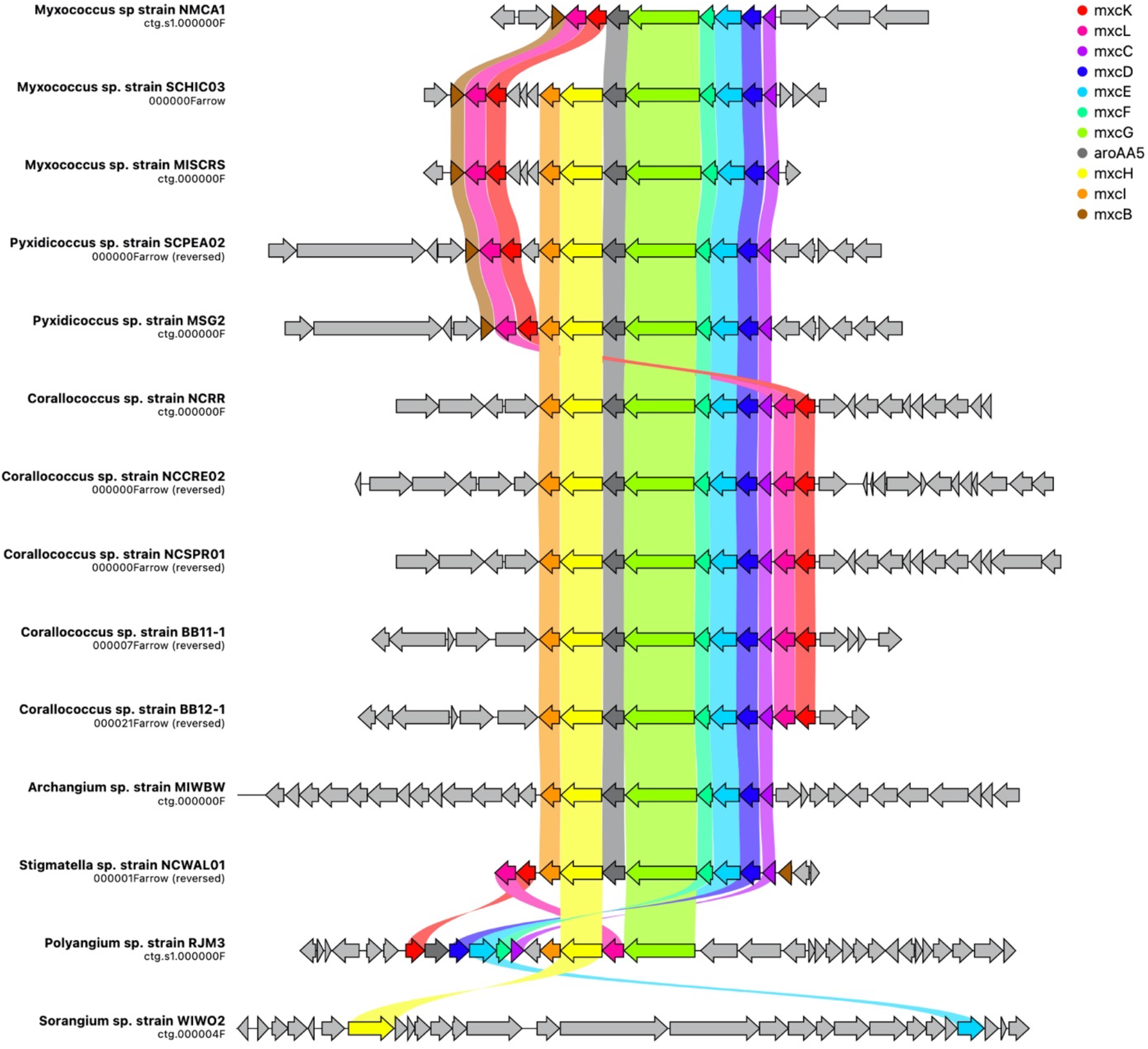
Spatial organization of the myxochelin BGC across all isolates with cluster homology. Gene names assigned by homology to the mxyochelin BGC deposited in the MiBIG database (BGC0001345), and ribbons indicate shared gene identity between clusters. Image generated using CAGECAT (version 1.0) using an identity threshold of 0.49 (94).

### Genomic organization of BGCs

AntiSMASH analysis of complete or near-complete genome data from FL3, MIWBW, MSG2, NCRR, NCSPR03, NMCA1, SCHIC03, and SCPEA02 provided contiguous sequence data sufficient to observe genome organization of BGCs. Cluster data from related myxobacteria with complete genome data from the antiSMASH database was used to compare BGC organization between related strains (Figures 6 and 7). Similarities between BGC content and genome organization were observed between subspecies (Figure 6A and 6B) and related strains within the same genus (Figures 6C and Figure 7). Biosynthetic gene clusters were dispersed throughout all genomes with segments of clusters appearing more dense with hybrid type clusters such as PKS-NRPS clusters or clusters including more than one cluster type. The myxochelin and myxoprincomide BGCs are located within a cluster-dense region in all sequenced environmental Myxococcia (Supplemental Figure 7). Clusters highly similar to carotenoid, geosmin, and VEPE/AEPE/TG-1 BGCs are often located in less-dense segments of analyzed chromosomes. Chromosomal segments with increased adjacency of hybrid and modular clusters were observed for all analyzed myxobacteria albeit less apparent in FL3 and *N. exedans* DSM 71^T^ (Figure 7C). Other than small differences in cluster content between analyzed subspecies, such as the absence of the myxovirescin BGC from NMCA1 (Figure 6A), numerous inversions of clusters resulting in changes to cluster organization were observed. Apparent BGC inversions were predominantly located within or near cluster-dense regions, and inversions were often relegated to core biosynthetic genes of single clusters with proximal genes unchanged between strains. Additional synteny analysis of strains with observed BGC inversions using a set of 10 homologous housekeeping genes revealed highly similar genome organization with no observed inversions (Supplemental Figures 8 and 9) (65).

**Figure 6:**
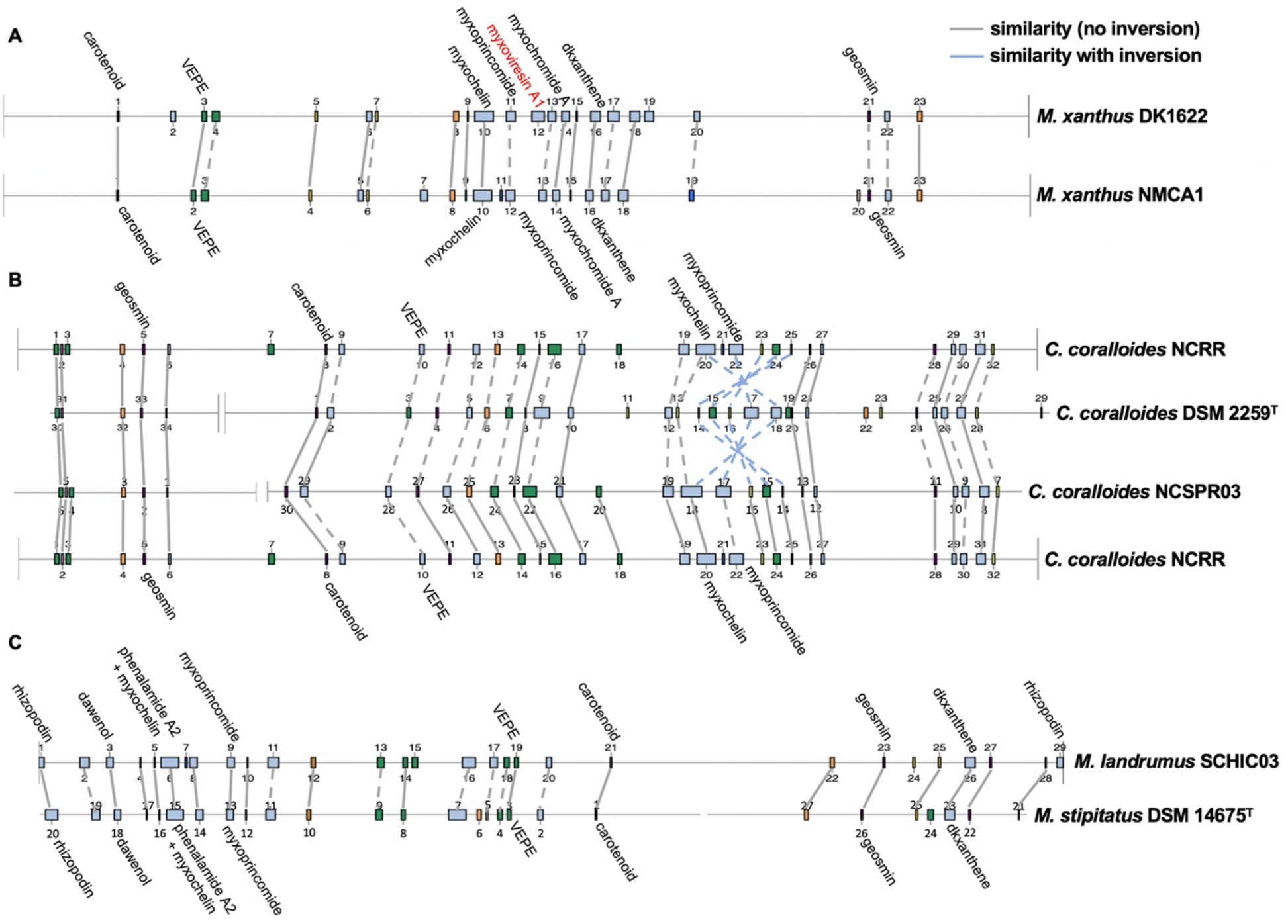
Whole genome comparisons depicting similar BGC organization between subspecies (A and B) and species (C). Labeled clusters have high similarity (≥66%) with corresponding myxobacterial BGCs deposited in the MIBiG database as determined by antiSMASH analysis. Myxobacterial clusters with no matching cluster between genomes are labeled red. Connecting lines between clusters indicate similar gene organization, and dashed lines indicate minor changes in gene organization or domain content of modular clusters. Inversions defined as inversion of entire clusters or subclusters within multicluster BGCs. BGC data for *M. xanthus* DK1622, *C. coralloides* DSM 2259^T^, and *M. stipitatus* DSM 14675^T^ obtained from the antiSMASH database (version 3). Numerical labels and graphical depictions of genome data taken from antiSMASH (version 6.0) with images altered to clearly depict similarities in cluster organization.

**Figure 7:**
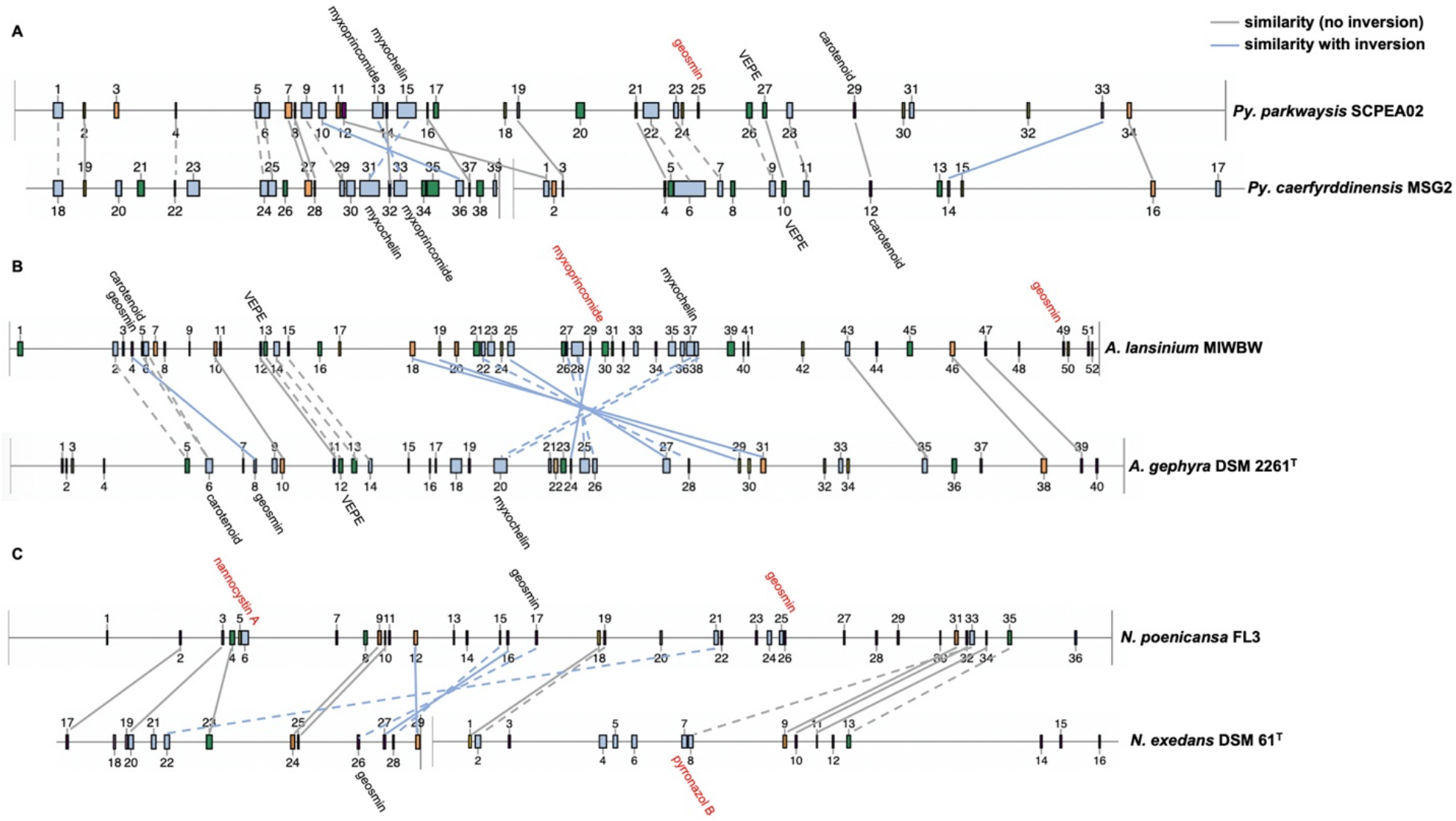
Cluster inversions impacting BGC organization between *Pyxidicoccus* (A), *Archangium* (B), and *Nannocystis* (C). Labeled clusters have high similarity (≥66%) with corresponding myxobacterial BGCs deposited in the MIBiG database as determined by antiSMASH analysis. Myxobacterial clusters with no matching cluster between genomes are labeled red. Connecting lines between clusters indicate similar gene organization, and dashed lines indicate slight changes in gene organization or domain content of modular clusters. Inversions defined as inversion of entire clusters or subclusters within multicluster BGCs. BGC data for *A. gephyra* DSM 2261^T^ and *N. exedans* DSM 61^T^ obtained from the antiSMASH database (version 3). Numerical labels and graphical depictions of genome data taken from antiSMASH (version 6.0) with images altered to clearly depict similarities in cluster organization.

### Proposal of nine novel species from seven genera of *Myxococcota*

We propose nine candidate strains to represent novel species with the genera *Archangium, Myxococcus, Nannocystis*, *Polyangium*, *Pyxidicoccus*, *Sorangium*, and *Stigmatella*. Comparative genomics including differences in genome content, phylogeny, and biosynthetic capacities as well as physiological and biochemical analyses support the following distinctions: *Archangium lansinium* sp. nov. (MIWBW^T^), *Myxococcus landrumus* sp. nov. (SCHIC03^T^), *Nannocystis bainbridgea* sp. nov. (BB15-2^T^), *Nannocystis poenicansa* sp. nov. (FL3^T^), *Nannocystis radixulma* sp. nov. (NCELM^T^), *Polyangium mundeleinium* sp. nov. (RJM3^T^), *Pyxidicoccus parkwaysis* sp. nov. (SCPEA02^T^), *Sorangium aterium* sp. nov. (WIWO2^T^), and *Stigmatella ashevillena* sp. nov. (NCWAL01^T^). Corresponding species descriptions for each candidate strain are provided below.

## DISCUSSION

### *Myxococcota* taxonomy

Myxobacteria are excellent resources for discovery of therapeutics and are suggested to be keystone taxa influencing polymicrobial community structure in soil (1,2, 66–68). Recent discoveries of novel *Corallococcus, Myxococcus*, and *Pyxidicoccus* species as well as species from lesser-studied genera indicate an abundance of uncharacterized myxobacteria (10–12, 14–17, 69–79). Our investigation of rhizospheric soil samples provided 20 environmental myxobacteria including 9 proposed novel species. As an initial attempt to isolate myxobacteria from soil, we were surprised by the effectiveness of morphology screening to enable discovery of myxobacteria from a variety of genera. We suspect that improved genome data will clarify the observed discrepancies in type strain differentiation and recommend high quality genome data be provided for all newly described type strain myxobacteria. We demonstrate that established comparative genome analysis thresholds for designation of novel species indicate *M. xanthus* DSM 16526^T^ and *M. virescens* DSM 2260^T^ to not be separate species as well as *St. aurantiaca* DSM 17044^T^, *Stigmatella erecta* DSM 16858^T^, and *Stigmatella hybrida* DSM 14722^T^. Data from sequenced *Myxococcus* and *Pyxidicoccus* strains aligns with previously recommended consideration of *Myxococcus/Pyxidicoccus* as a single genus (11, 12). However, we note the significant differences in BGC content between *Myxococcus* and *Pyxidicoccus* strains included in this analysis. Overall, our phylogenetic analysis provides further support for comparative genomics approaches to identify and classify myxobacteria. The primary limitation being the absence of quality genome data for established type strains within genera such as *Archangium, Polyangium*, and *Sorangium*.

### Expansion of the genus *Nannocystis*

Current type strain *Nannocystis* include *N. exedans* DSM 71^T^, *N. konarekensis* DSM 104509^T^, and *N. pusilla* DSM 14622^T^. Our investigation resulted in discovery of an additional three proposed type strains doubling the current total of *Nannocystis* (17). We also report genome data for all three proposed type strain species as well as three *Nannocystis* subspecies and *N. pusilla* DSM 14622^T^. Cumulative analysis of all sequenced *Nannocystis* and our isolates provided notable differences in BGC content when compared to other myxobacteria such as absence of a cluster with similarity to the myxochelin BGC, presence of multiple siderophore and phosphonate cluster types, and smaller cluster sizes. Resulting genome data for an additional eight *Nannocystis* will improve future efforts to characterize and describe members of this underexplored genus of myxobacteria.

### Genomic organization of BGCs and adaptability of specialized metabolism

Afforded complete or near-complete genome sequence data, we report the first comparative analysis of spatial organization of myxobacterial BGCs. Dissimilar from *Streptomyces* localization of BGCs in the extremities of linear genomes (80, 81), clusters are distributed throughout the circular genomes of myxobacteria. Observed cluster-rich regions replete with modular BGCs and noted inversions of biosynthetic genes could however contribute to metabolic differentiation similar to how terminal compartments of *Streptomyces* chromosomes enable spatial reorganization and conditional expression of BGCs during metabolic development and sporulation (80). We suggest BGC enriched regions may benefit BGC evolution and contribute to metabolic adaptability of myxobacteria. Compartmentalization of modular-type clusters with highly homologous domains may benefit module duplication and deletion events associated with evolution of BGCs (82–84). Vertical inheritance of the myxochelin cluster is apparent from all sequenced Myxococcia. Our data also reveals vertical transfer and likely concerted evolution of myxoprincomide-type clusters across sequenced Myxococcia (84). Alternatively, presence of the myxovirescin trans-AT PKS cluster within a cluster-rich region of the *M. xanthus* DK1622 genome and absence of a homologous cluster in NMCA1 indicates horizontal acquisition. Although loss of the myxovirescin cluster from NMCA1 provides an alternative explanation, absence of the myxovirescin cluster from all other sequenced members of the *Myxococcus* and Myxococcota currently deposited in the antiSMASH database instead supports horizontal acquisition by DK1622. Regardless, either explanation for cluster presence in DK1622 and absence in other myxobacteria demonstrates metabolic adaptability within the BGC enriched segments of myxobacterial genomes. Further investigation of chromosomal organization of BGCs in myxobacteria is required to determine functional impacts on metabolic adaptability and cluster evolution.

### Species Descriptions

#### (i) *Archangium lansinium* sp. nov

*Archangium lansinium* (lan.sin’i.um. N.L. neut. ajd. *lansinium* from Lansing, Michigan, United States of America, referencing the area of isolation).

Vegetative cells glide on solid media. Cells grow as translucent swarms during the early growth phase on VY/4 agar and form non-uniform, clumping fruiting bodies that range from yellow to pale orange over time. Aerobic growth was observed at 20 to 35°C but not at 40°C at a pH range of 6.0 to 8.0. Hydrolyzes gelatin and esculin. Shows an API ZYM positive reaction to alkaline phosphatase, C4 esterase, C8 lipase, C14 lipase, leucine arylamidase, valine arylamidase, cysteine arylamidase, trypsin, α-chymotrypsin, acid phosphatase, napthol-AS-BI-aphosphohydrolase, and *N*-acetyl-β-glucosaminidase and a negative reaction to α-galactosidase, β-galactosidase, β-glucuronidase, α-glucosidase, β-glucosidase, α-mannosidase, and α-fucosidase. DNA GC content is 68.1%. The genome assembly of the organism is available at NCBI Assembly (ASM2662663v1). The 16S ribosomal RNA gene sequence is available at Genbank (OP852336.1). Phylogenetically most similar to *A. gephyra* DSM 2261^T^.

The type strain (MIWBW^T^ = NCCB 100916^T^) was isolated from soil collected in the Summer of 2021 from the roots of a white basswood tree near the city of Lansing, Michigan, United States of America (42.73°N, 84.48°W).

#### (ii) *Myxococcus landrumus* sp. nov

*Myxococcus landrumus* (lan.drum’us. N.L. masc. ajd. *landrumus* from Landrum, South Carolina, United States of America, referencing the area of isolation).

Vegetative cells glide on solid media. Cells grow as slightly orange swarms and develop rounded, stalking fruiting bodies that range from white to purple on VY/4 media. Aerobic growth was observed at 20 to 35°C but not at 40°C at a pH range of 6.0 to 10.0. Hydrolyzes gelatin and esculin. Shows an API ZYM positive reaction to C4 esterase, C8 lipase, C14 lipase, leucine arylamidase, valine arylamidase, acid phosphatase, and napthol-AS-BI-aphosphohydrolase and a negative reaction to alkaline phosphatase, cysteine arylamidase, trypsin, α-chymotrypsin, *N*-acetyl-β-glucosaminidase, α-galactosidase, β-galactosidase, β-glucuronidase, α-glucosidase, β-glucosidase, α-mannosidase, and α-fucosidase. DNA GC content is 68.6%. The genome assembly of the organism is available at NCBI Assembly (ASM1730163v1). The 16S ribosomal RNA gene sequence is available at Genbank (OP852328v1). Phylogenetically most similar to *M. stipitatus* DSM 14675^T^.

The type strain (SCHIC03^T^ = NCCB 100915^T^) was isolated from soil collected in Spring 2020 from the roots of a hickory tree near the city of Landrum, South Carolina, United States of America (35.14°N, −82.16°W).

#### (iii) *Nannocystis bainbridgea* sp. nov

*Nannocystis bainbridgea* (bain’bridg.ea. N.L. fem. ajd. *bainbridgea* from Bainbridge Island, Washington, United States of America, referencing the area of isolation).

Vegetative cells glide on solid media. Cells are agarolytic and form deep etches in VY/4 agar. Grows as translucent red swarms and forms rounded fruiting bodies. Aerobic growth was observed at 20 to 30°C but not at 35 to 40°C at a pH range of 7.0 to 8.0. Hydrolyzes gelatin and esculin. Shows an API ZYM positive reaction to alkaline phosphatase, C4 esterase, C8 lipase, leucine arylamidase, α-chymotrypsin, acid phosphatase, and napthol-AS-BI-aphosphohydrolase and a negative reaction to C14 lipase, valine arylamidase, cysteine arylamidase, trypsin, *N*-acetyl-β-glucosaminidase, α-galactosidase, β-galactosidase, β-glucuronidase, α-glucosidase, β-glucosidase, α-mannosidase, and α-fucosidase. DNA GC content is 71.5%. The genome assembly of the organism is available at NCBI Assembly (ASM2696555v1). The 16S ribosomal RNA gene sequence is available at Genbank (OP852329.1). Phylogenetically most similar to *N. exedens* DSM 71^T^.

The type strain (BB15-2^T^) was isolated from soil collected summer of 2020 from the roots of a blueberry bush near the city of Bainbridge Island, Washington, United States of America (47.65°N, −122.55°W).

#### (iv) *Nannocystis poenicansa* sp. nov

*Nannocystis poenicansa* (poe.ni.can’sa. N.L. fem. adj. *poenicansa* the bright red, referencing bright red pigment production).

Vegetative cells glide on solid media. Cells are agarolytic and form deep etches in VY/4 agar. Aerobic growth was observed at 20 to 35°C but not at 40°C at a pH range of 6.0 to 8.0. Hydrolyzes esculin. Shows an API ZYM positive reaction to alkaline phosphatase, C4 esterase, C8 lipase, C14 lipase, leucine arylamidase, cysteine arylamidase, acid phosphatase, napthol-AS-BI-aphosphohydrolase, and β-glucosidase and a negative reaction to valine arylamidase, trypsin, α-chymotrypsin, *N*-acetyl-β-glucosaminidase, α-galactosidase, β-galactosidase, β-glucuronidase, α-glucosidase, α-mannosidase, and α-fucosidase. DNA GC content is 71.5%. The complete genome sequence of the organism is available at GenBank (CP114040.1). The 16S ribosomal RNA gene sequence is available at Genbank (OP852330.1). Phylogenetically most similar to *N. exedens* DSM 71^T^.

The type strain (FL3^T^ = NCCB 100918^T^) was isolated from soil collected Fall of 2020 from the roots of a southern live oak near the city of Palm Coast, Florida, United States of America (29.59°N, −81.21°W).

#### (v) *Nannocystis radixulma* sp. nov

*Nannocystis radixulma* (ra’dix.ul.ma. L. fem. n. *poenicansa* the root of elm, referencing the isolation from an elm tree rhizosphere).

Vegetative cells glide on solid media. Cells are agarolytic and form deep etches in VY/4 agar. Early growth ranges from translucent swarm perimeters to yellow pigmented swarm centers, and form textured, clumping fruiting bodies that range from yellow to orange on VY/4 agar. Aerobic growth was observed at 25 to 30°C but not at 20°C or 35 to 40°C at a pH range of 6.0 to 9.0. Hydrolyzes gelatin and esculin. Shows an API ZYM positive reaction to alkaline phosphatase, C4 esterase, C8 lipase, leucine arylamidase, cysteine arylamidase, acid phosphatase, napthol-AS-BI-aphosphohydrolase, and β-glucosidase and a negative reaction to C14 lipase, valine arylamidase, trypsin, α-chymotrypsin, *N*-acetyl-β-glucosaminidase, α-galactosidase, β-galactosidase, β-glucuronidase, α-glucosidase, α-mannosidase, and α-fucosidase. DNA GC content is 71.2%. The genome assembly of the organism is available at NCBI Assembly (ASM2836909v1). The 16S ribosomal RNA gene sequence is available at Genbank (OP852331.1). Phylogenetically most similar to *N. exedens* DSM 71^T^.

The type strain (NCELM^T^ = NCCB 100919^T^) was isolated from soil collected in Spring 2020 from the roots of an elm tree near the city of Asheville, North Carolina, United States of America (35.63°N, −82.55°W).

#### (vi) *Polyangium mundeleinium* sp. nov

*Polyangium mundelenium* (mun.del.en’ni.um. N.L. neut. adj. *mundelenium* from Mundelein, Illinois, United States of America, referencing the area of isolation).

Vegetative cells glide on solid media. Cells grow as translucent swarms and develop dispersed rounded, yellow fruiting bodies on VY/4 media. Aerobic growth was observed at 20 to 30°C but not at 20°C or 35 to 40°C at a pH of 7.0. Hydrolyzes gelatin and esculin. Shows an API ZYM positive reaction to alkaline phosphatase, C4 esterase, C8 lipase, leucine arylamidase, valine arylamidase, α-chymotrypsin, acid phosphatase, and napthol-AS-BI-aphosphohydrolase and a negative reaction to C14 lipase, cysteine arylamidase, trypsin, *N*-acetyl-β-glucosaminidase, α-galactosidase, β-galactosidase, β-glucuronidase, α-glucosidase, β-glucosidase, α-mannosidase, and α-fucosidase. DNA GC content is 68.6%. The genome assembly of the organism is available at NCBI Assembly (ASM2836910v1). The 16S ribosomal RNA gene sequence is available at Genbank (OP852332.1). Phylogenetically most similar to *P. fumosum* DSM 14688^T^.

The type strain (RJM3^T^ = NCCB 100920^T^) was isolated from soil collected Fall of 2020 from the roots of a red Japanese maple near the village of Mundelein, Illinois, United States of America (42.26°N, −88.0°W).

#### (vii) *Pyxidicoccus parkwaysis* sp. nov

*Pyxidicoccus parkwaysis* (park.way.sis. N.L. masc. adj. *parkwaysis* from Parkway Farm in Landrum, South Carolina, United States of America, referencing the area of isolation).

Vegetative cells glide on solid media. Cells grow as mucoid swarms with a slight pink pigmentation and develop, mounded fruiting bodies after 3 weeks of growth on VY/4 media. Aerobic growth was observed at 20 to 40°C at a pH range of 6.0 to 9.0. Hydrolyzes gelatin and esculin. Shows an API ZYM positive reaction to C4 esterase, C8 lipase, leucine arylamidase, acid phosphatase, and napthol-AS-BI-aphosphohydrolase and a negative reaction to alkaline phosphatase, C14 lipase, valine arylamidase, cysteine arylamidase, trypsin, α-chymotrypsin, *N*-acetyl-β-glucosaminidase, α-galactosidase, β-galactosidase, β-glucuronidase, α-glucosidase, β-glucosidase, α-mannosidase, and α-fucosidase. DNA GC content is 69.6%. The genome assembly of the organism is available at NCBI Assembly (ASM1730173v1). The 16S ribosomal RNA gene sequence is available at Genbank (OP852333). Phylogenetically most similar to *P. caerfyrddinensis* CA032A^T^.

The type strain (SCPEA02^T^ = NCCB 100921^T^) was isolated from soil collected in Spring 2020 from the roots of a peach tree near Parkway Farm in Landrum, South Carolina, United States of America (35.14°N, −82.12°W).

#### (viii) *Sorangium aterium* sp. nov

*Sorangium aterium* (at’er.i.um. N.L. neut. adj. *aterium* the flat black, referencing black pigment production).

Vegetative cells are Gram negative and glide on solid media. Degrades and decomposes cellulose filter paper. Cells are pigmented dark brown to black when grown on VY/4 agar. Dark brown to black fruiting bodies form on ST21 media, and similarly pigmented sporangioles from on VY/4 agar. Aerobic growth was observed at 25 to 30°C but not at 20°C or 35 to 40°C at a pH range of 6.0 to 7.0. Hydrolyzes esculin. Shows an API ZYM positive reaction to alkaline phosphatase, C4 esterase, C8 lipase, leucine arylamidase, acid phosphatase, napthol-AS-BI-aphosphohydrolase, and β-glucosidase and a negative reaction to C14 lipase, valine arylamidase, cysteine arylamidase, trypsin, α-chymotrypsin, *N*-acetyl-β-glucosaminidase, α-galactosidase, β-galactosidase, β-glucuronidase, α-glucosidase, α-mannosidase, and α-fucosidase. DNA GC content is 71.1%. The genome assembly of the organism is available at NCBI Assembly (ASM2836893v1). The 16S ribosomal RNA gene sequence is available at Genbank (OP852334.1). Phylogenetically most similar to *S. cellulosum* Soce56.

The type strain (WIWO2^T^ = NCCB 100922^T^) was isolated from soil collected in Fall 2021 from the roots of a white oak near the village of Pleasant Prairie, Wisconsin, United States of America (42.56°N, −87.94°W).

#### (ix) *Stigmatella ashevillena* sp. nov

*Stigmatella ashvillena* (ash’vill.en.a. N.L. fem. adj. from Asheville, North Carolina, United States of America, referencing the area of isolation).

Vegetative cells are Gram negative and glide on solid media. Cells are yellow during early growth on VY/4 media and form stalked, orange fruiting bodies over time. Aerobic growth was observed at 25 to 30 °C but not at 20°C or 35 to 40°C at a pH range of 6.0 to 9.0. Hydrolyzes gelatin and esculin. Shows an API ZYM positive reaction to alkaline phosphatase, C4 esterase, C8 lipase, C14 lipase, leucine arylamidase, cysteine arylamidase, acid phosphatase, napthol-AS-BI-aphosphohydrolase, α-glucosidase, β-glucosidase, and *N*-acetyl-β-glucosaminidase and a negative reaction to valine arylamidase, trypsin, α-chymotrypsin, α-galactosidase, β-galactosidase, β-glucuronidase, α-mannosidase, and α-fucosidase. DNA GC content is 66.9%. The genome assembly of the organism is available at NCBI Assembly (ASM2836897v1). The 16S ribosomal RNA gene sequence is available at Genbank (OP852335.1). Phylogenetically most similar to *S. aurantiaca* DSM17044^T^.

The type strain (NCWAL01^T^ = NCCB 100923^T^) was isolated from soil collected in Spring 2020 from the roots of a walnut tree near the city of Asheville, North Carolina, United States of America (35.63°N, −82.55°W).

## MATERIALS AND METHODS

### Isolation of myxobacteria

Bacteriolytic myxobacteria were isolated by the *Escherichia coli* baiting method (18). Briefly, an *E. coli* lawn was grown overnight at 37°C and resuspended in 1 mL of an antifungal solution (250 μg/mL cycloheximide and nystatin). A 300 μl solution was spread across a WAT agar (1.5% agar, 0.1% CaCl_2_) plate and air dried. Previously air dried soil was wet with the antifungal solution to a mud like consistency, and a pea sized amount was placed on the dried *E. coli* WAT plate. The plate was incubated at 25°C for up to 4 weeks. After 3 days of incubation, the plates were checked daily for the appearance of lytic zones or fruiting bodies in the *E. coli* lawn. Using a syringe needle, the lytic zones were moved to a plate of VY/4 (Baker’s yeast 2.5 g/L, CaCl_2_ x 2H_2_O 1.36 g/L, vitamin B12 0.5 mg/L, agar 15 g/L). The swarm edge was repeatedly used to inoculate a fresh VY/4 plate until pure cultures were obtained. Isolates were cultivated continuously at 25-30°C on VY/4. Isolation of cellulolytic myxobacteria was accomplished using the filter paper method (18). A small square of autoclaved filter paper was placed in the center of a ST21 agar plate (K_2_HPO_4_ 1 g/L, yeast extract 20 mg/L, agar 14 g/L, KNO_3_ 1 g/L, MgSO_4_ x 7H_2_O 1 g/L, CaCl_2_ x 2H_2_O 1 g/L, MnSO_4_ x 7H_2_O 0.1 g/L, FeCl_3_ 0.2 g/L). A pea sized amount of soil, wet with the antifungal solution, was placed at the edge of the filter paper. Plates were incubated at 25°C for up to 2 months. After 2 weeks, the plates were checked every two days for cellulose degradation and fruiting body formation. Fruiting bodies were moved to a fresh ST21 plate with filter paper repeatedly until pure monocultures were observed. Isolates were cultivated continuously at 25-30°C on VY/4.

### Cultivation of isolates

All isolates were maintained on VY/4 media. Growth in liquid cultures was achieved using CYH/2 media (casitone 0.75 g/L, yeast extract 0.75 g/L, starch 2g/L, soy flour 0.5 g/L, glucose 0.5 g/L, MgSO_4_ x 7H_2_O 0.5 g/L, CaCl_2_ x 2H2O 1 g/L, HEPES 6 g/L, EDTA-Fe 8 mg/L, vitamin B12 0.5 mg/L)

### Sequencing methods

Once pure cultures were obtained, DNA isolation was performed using either the Monarch HMW DNA extraction kit for tissue or the Macherey Nagel nucleobond HMW DNA extraction kit, following the manufacturer’s instructions for Gram-negative bacteria. All novel strains were sequenced using Pacbio Sequel or Sequel II with a 10 hour movie. De Novo Assembly of the genome was accomplished using the SMRT Analysis Hierarchical Genome Assembly Process (HGAP; SMRT Link 9.0.0 or SMRT Link 10.1.0). *Nannocystis pusilla* Na p29^T^ was sequenced using an Oxford Nanopore Minion Flongle R9.4.1. Raw data was processed using Guppy (v6.0.1) (85). Reads were trimmed using porechop (v0.2.4) (86). The worst 10% of reads were filtered out using filtong (v0.2.1). Flye assembler (v2.9) was used for de novo assembly using the trimmed and filtered reads (87). Correction of the final assembly was achieved by long read correction using two iterations of Racon (v1.5) (88), followed by two iterations of Medaka (v1.7.2) (89). Genome assemblies and complete genomes data was deposited at NCBI for the following isolates: *Corallococcus exiguus* strain NCCRE02 (ASM1730297v1), *Corallococcus* sp. strain NCSPR01 (ASM1730913v1), *Corallococcus* sp. strain BB11-1 (ASM2662662v1), *Corallococcus* sp. strain BB12-1 (ASM2662676v1), *Corallococcus* sp. strain NCRR (ASM2696553v1), *Myxococcus* sp. strain MISCRS (ASM2662660v1), *Myxococcus* sp. strain NMCA1 (ASM2681020v1), *Nannocystis* sp. strain SCPEA4 (ASM2662668v1), *Nannocystis* sp. strain ILAH1 (ASM2662658v1), *Nannocystis* sp. strain RBIL2 (ASM2662674v1) and *Pyxidicoccus* sp. strain MSG2 (ASM2662670v1). Genome assembly for *Nannocystis pusilla* Na p29^T^ was also deposited at NCBI (ASM2662666v1).

### Microscopy

A Zeiss stereo discovery.V12 microscope using Axio cam 105 and a Plano Apo S 1.0X objective were used to observe fruiting bodies and swarming patterns.

### Comparative genomics

OrthoAni calculations and tree generation were achieved using OAT (orthoANI tool v0.93.1) (90). dDDH calculations were performed on the type strain genome server (TYGS) website (91). Synteny analysis was done using SimpleSynteny (v1.6) (65).

### Enzymatic Assays

Enzymatic activity was assessed for myxobacteria utilizing commercial API ZYM (bioMérieux, France) and API NE (bioMérieux, France) kits. Each isolate strain was suspended in NaCl 0.85% to an OD_600_ of 0.7 and 0.1 for API NE. API ZYM strips were incubated for 4.5 hours at 37°C, and API NE strips were incubated for 24 hours at 37°C. After incubation, specific reagents were added to the cupule and evaluated according to the manufacturer’s instructions.

### Growth Conditions

For most of the myxobacteria tested, strains were grown on VY/4 (pH 7.2) for 5 to 7 days and resuspended in deionized water to an OD_600_ of 0.5. For the genera *Nannocystis* and *Polyangium*, strains were grown for 5 to 7 days in CYH/2 media, centrifuged, washed, and resuspended in sterile distilled water. The optimal growth temperature was tested by inoculating VY/4 plates with 25 μl of the 0.5 OD_600_ suspension for given myxobacteria. Plates were incubated at 20, 25, 30, 35, and 40°C for up to 14 days. Optimal pH was assessed by plating the 0.5 OD_600_ solution on VY/4 plates buffered to pH 5, 6, 7, 8, 9, or 10. The pH conditions were buffered at a pH of 5 to 6 with 25 mM MES buffer, 7 to 8 with 25 mM HEPES buffer, and 9 to 10 with 25 mM TRIS buffer in VY/4 plates and incubated at 25°C for two weeks. Comparisons of swarm diameters were used to determine optimal growth conditions.

### BGC analysis

FASTA files for all sequenced isolates were uploaded for analysis using antiSMASH (version 6.1.1) using relaxed detection strictness with all extra features toggled on (28). Resulting antiSMASH job IDs from analyzed isolates were submitted as queries using the BIGFAM database v1.0.0 (1,225,071 BGCs and 29,955 GCFs) to assess BGC similarity to database clusters (31). All BGCs with >900 distance from model GCFs were subsequently dereplicated manually to remove characterized BGCs not clustered with GCGs within the BIGFAM database. The antiSMASH database v3.0 (147,571 BGCs) was used to analyze BGCs from included myxobacteria (92). BiG-SCAPE v1.1.0 was used to analyze all BGCs .gbk files from sequenced isolates as well as all .gbk files from myxobacteria with sequenced genomes deposited in the antiSMASH database with the using the “hybrids-off” and “MiBIG” parameters (33, 93).

### Supplementary Materials

Supplementary materials include: dDDH values for all isolates (Supplementary Table 1), a map of collection sites for all isolates (Supplementary Figure 1), a phylogenetic tree for isolates and type strain myxobacteria from 16S RNA sequences (Supplementary Figure 2), orthoANI values for all isolates not included in Figure 2 (Supplementary Figure 3), BGC similarity networks generated with BiG-SCAPE (Supplementary Figures 4 and 5), chromosomal context for the myxovirescin BGC in *M. xanthus* DK1622 and NMCA1 (Supplementary Figure 6), evolutionary relationship the myxochelin BGC (Supplementary Figure 7), and shared synteny of myxobacteria included in Figures 6 and 7 (Supplementary Figures 8 and 9).

## Author Contributions

Isolation of environmental myxobacteria A.A., K.E.P., and T.K.; growth profiles, biochemical assays, and imaging for isolates K.E.P., T.K., and M.H.; genome sequencing A.A., K.E.P., and S.E.D.; BGC analyses A.A., K.E.P., and D.C.S.; manuscript preparation and editing A.A., K.E.P., and D.C.S.; supervision and administration D.C.S. All authors have read and approved the final manuscript.

## Acknowledgements

The authors would like to thank Mary Williams and Ashlen Ahearne for coordinating the collection and delivery of soil samples analyzed in this project. The authors also appreciate the Glycoscience Center of Research Excellence Imaging Research Core and Computational Chemistry and Bioinformatics Research Core for assistance imaging isolates and assembly of the *N. pusilla* Na p29^T^ draft genome with specific gratitude for the Imaging Research Core manager, Dr. Ruofan Cao.

