## Supplemental Table and Figures for "Chromosomal organization of biosynthetic gene clusters suggests plasticity of myxobacterial specialized metabolism including descriptions for nine novel species: *Archangium lansinium* sp. nov., *Myxococcus landrumus* sp. nov., *Nannocystis bainbridgea* sp. nov., *Nannocystis poenicansa* sp. nov., *"

### Supplemental Figures and Tables

**Supplemental Table 1:** dDDH values between isolates and closest relatives. All named myxobacteria are type strains excluding *Stigmatella aurantiaca* DW4-1 and *Sorangium cellulosum* Soce56. The established cutoff for new species designation is <70%.

| *Isolate* | *Closest relative(s) dDDH* |
|---|---|
| **BB15-2** | *Nannocystis exedens* 36.2% |
| **NCELM1** | FL3 33.2%<br>*Nannocystis exedens* 33.1% |
| **RBIL2** | *Nannocystis pusilla* 55.2% |
| **FL3** | *Nannocystis exedens* 40.4% |
| **SCHIC03** | *Myxococcus stipitatus* 49.4% |
| **MIWBW** | *Archangium violaceum* 34.7% |
| **RJM3** | *Polyangium fumosum* 55.6% |
| **SCPEA02** | *Pyxidicoccus caerfyrddinensis* 34.4% |
| **NCWAL1** | *Stigmatella aurantiaca* DW4-1 43.2% |
| **WIWO2** | *Sorangium cellulosum* Soce56 56.6% |
| **NCSPR01** | *Corallococcus corraloides* 54% |
| **NCRR** | NCSPR01 100% |
| **NCCRE02** | *Corallococcus exiguus* 66.7% |
| **BB11-1** | *Corallococcus soli* 87% |
| **BB12-1** | *Corallococcus terminator* 65.5% |
| **MSG2** | *Pyxidicoccus caerfyrddinensis* 54.1% |
| **MISCRS** | *Myxococcus fulvus* 63% |
| **NMCA1** | *Myxococcus virescens* 65% |
| **SCPEA4** | NCELM 70.8% |
| **ILAH1** | *Nannocystis pusilla* 55.1% |

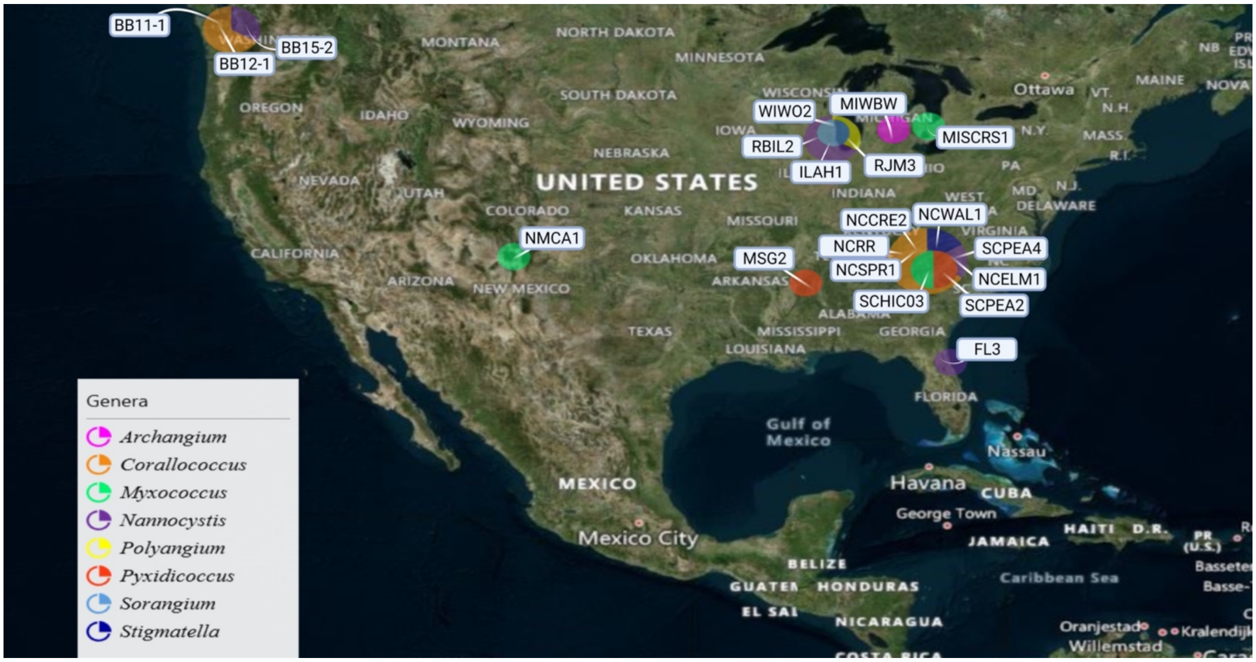


**Supplemental Figure 1:** Map of collection sites for all discussed myxobacterial isolates.

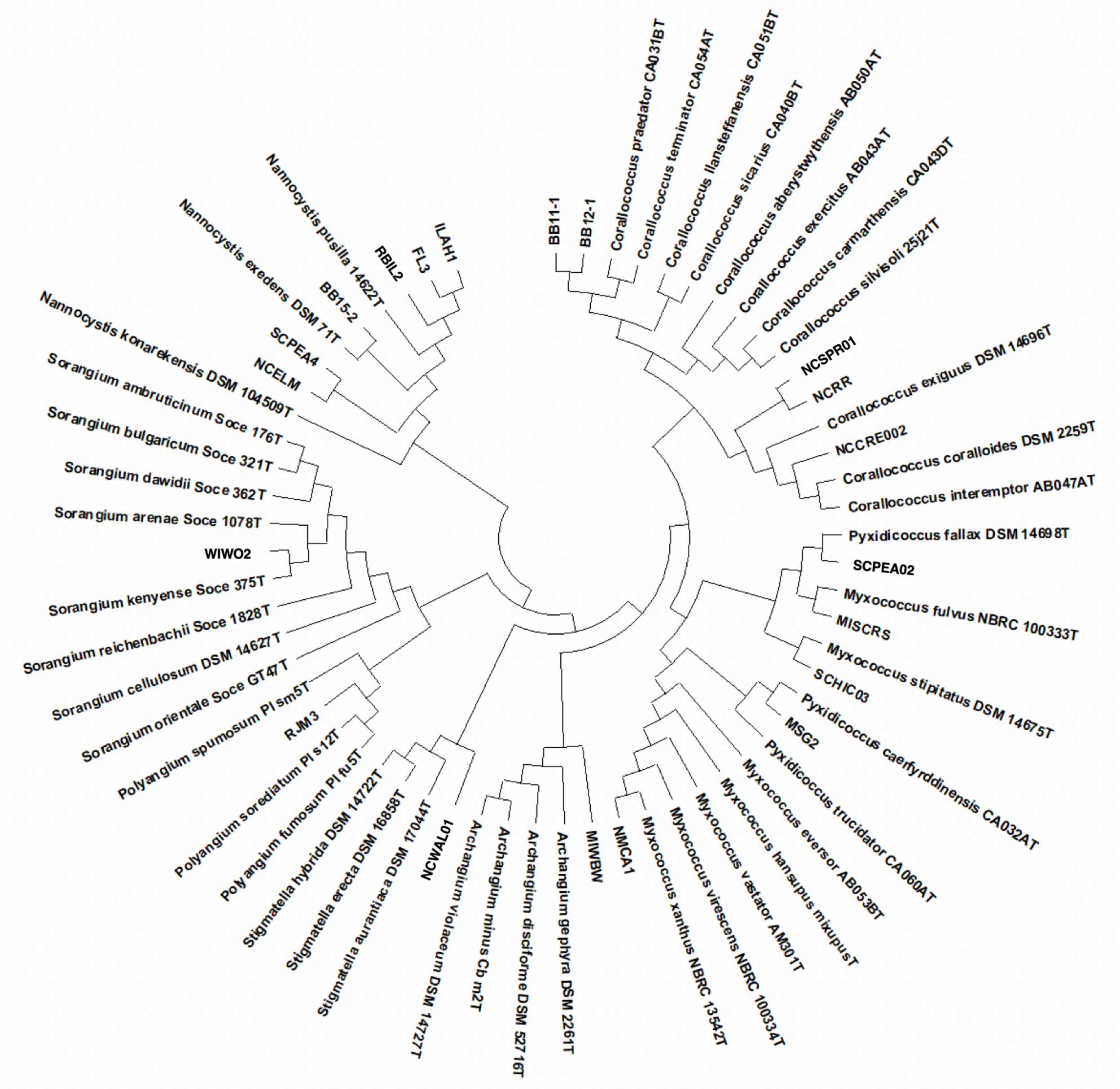


**Supplemental Figure 2:** Nearest neighbor phylogenetic tree from 16S RNA sequences comparing myxobacterial isolates with type strain myxobacteria.

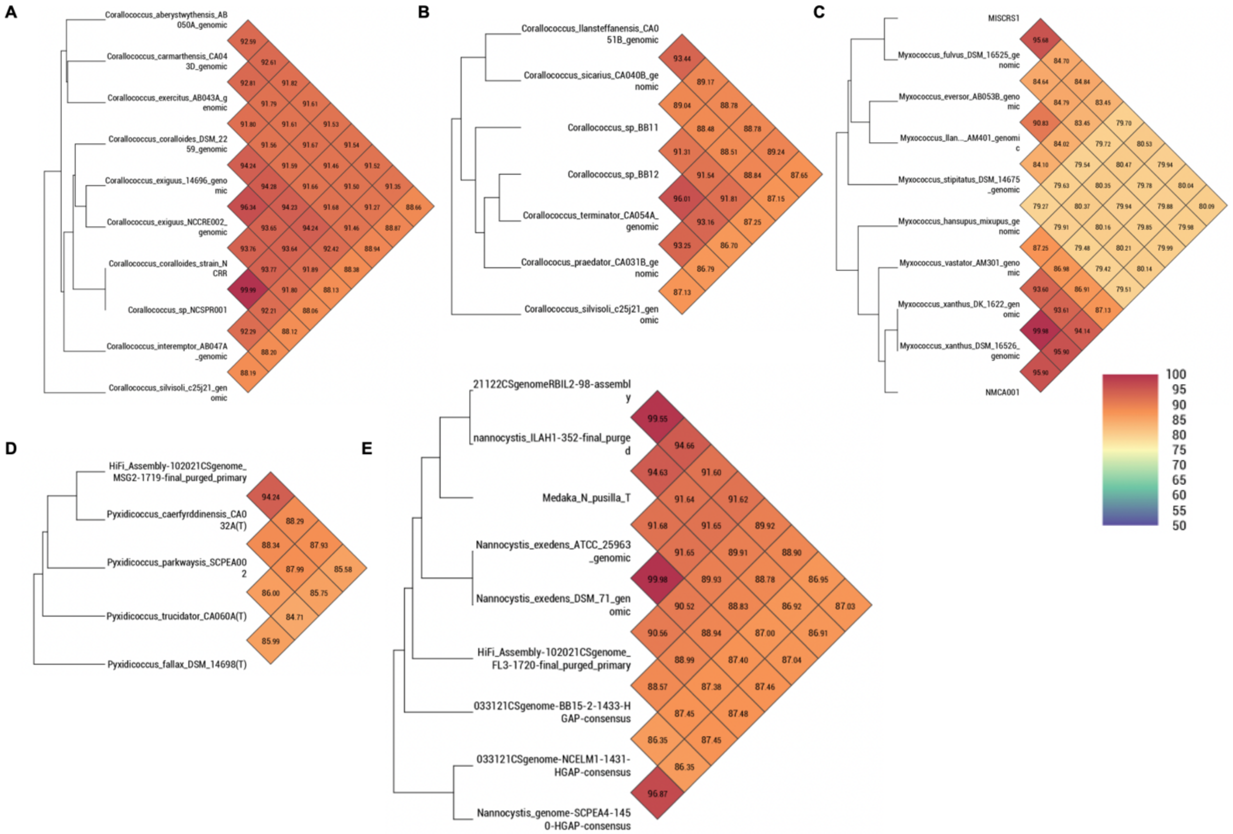


**Supplemental Figure 3:** Heatmaps generated from OrthoANI values calculated using OAT for A) *Corallococcus* spp. including isolates NCCRE02, NCRR, and NCSPR01; B) *Corallococcus* spp. including isolates BB11-1 and BB12-1; C) *Myxococcus* spp. including isolates MISCRS1 and NMCA1; D) *Pyxidicoccus* spp. including isolates MSG2 and SCPEA02; E) *Nannocystis* spp. including isolates RBIL2, ILAH1, FL3, BB15-2, NCELM, and SCPEA4.

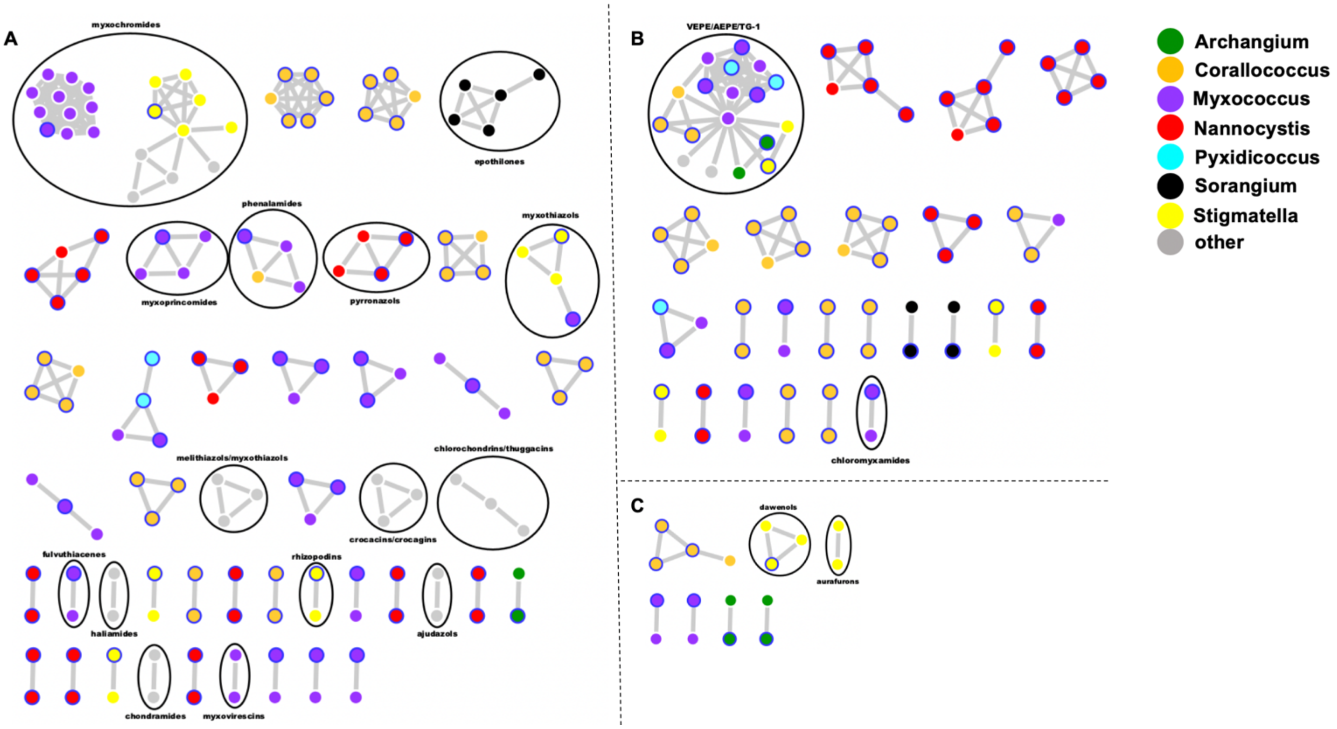


**Supplemental Figure 4:** BGC similarity network generated with BiG-SCAPE (v1.1.0) depicting cluster similarity between BGCs from myxobacterial isolates and sequenced myxobacteria in the antiSMASH database and cluster types determined by antiSMASH analysis. A) PKS-NRPS hybrid BGCs, B) NRPS BGCs, C) type I PKS BGCs. Gene cluster families that include a characterized BGC deposited in the BGC database are circled with the cluster product provided as text.

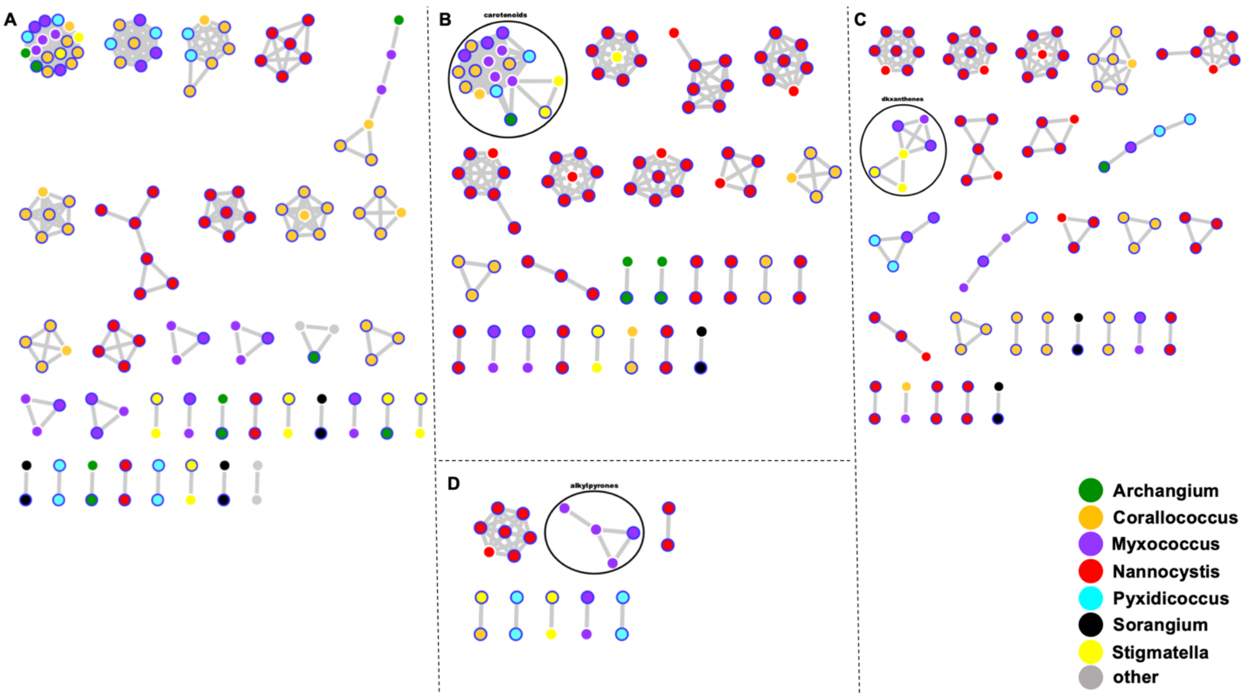


**Supplemental Figure 5:** BGC similarity network generated with BiG-SCAPE (v1.1.0) depicting cluster similarity between BGCs from myxobacterial isolates and sequenced myxobacteria in the antiSMASH database and cluster types determined by antiSMASH analysis. A) RiPPs BGCs, B) terpene BGCs, C) others BGCs, D) PKS (other) BGCs. Gene cluster families that include a characterized BGC deposited in the BGC database are circled with the cluster product provided as text.

A

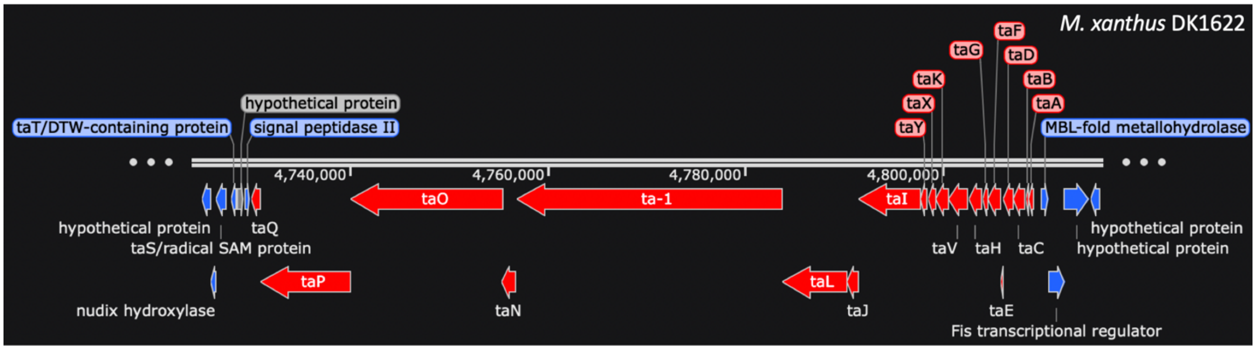


B

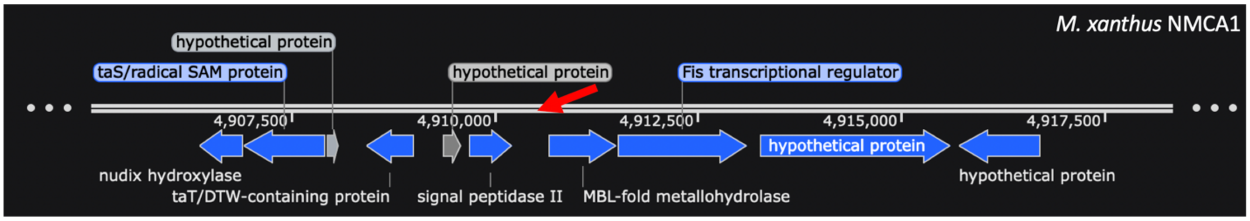


**Supplemental Figure 6:** Chromosomal context for A) the myxovirescin BGC from *M. xanthus* DK1622 and B) the same region for *M. xanthus* NMCA1 missing the myxovirescin BGC. Homologous features in blue, missing myxovirescin BGC features in red, and the red arrow indicates the missing myxovirescin BGC in *M. xanthus* NMCA1.

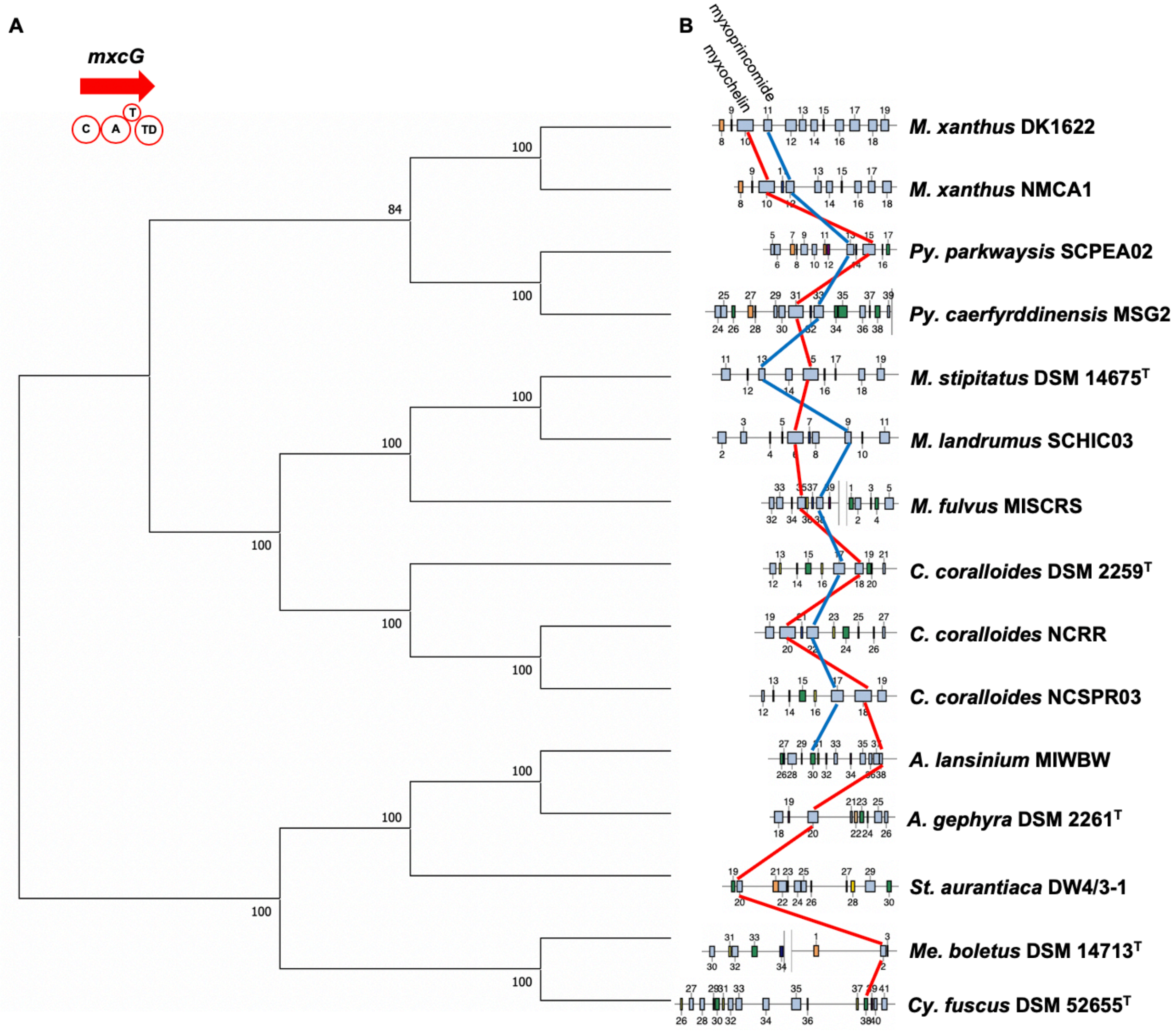


**Supplemental Figure 7:** Evolutionary relationships of the myxochelin cluster. Evolutionary history of *mxcG* using the Minimum Evolution method with percentage of replicate trees including taxa as depicted in a 300 replicate bootstrap test done in MEGA X (A). High cluster density genome regions tracking myxochelin (red) and myxoprincomide (blue) BGCs for analyzed myxobacteria included as branch labels (B). BGC data for *M. xanthus* DK1622, *M. stipitatus* DSM 14675$^T$, *C. coralloides* DSM 2259$^T$, *A. gephyra* DSM 2261$^T$, *St. aurantiaca* DW4/3-1, *Me. boletus* DSM 14713$^T$, and *Cy. fuscus* DSM 52655$^T$ obtained from the antiSMASH database (version 3). Numerical labels and graphical depictions of genome data taken from antiSMASH (version 6.0) with images altered to clearly depict similarities in cluster organization.

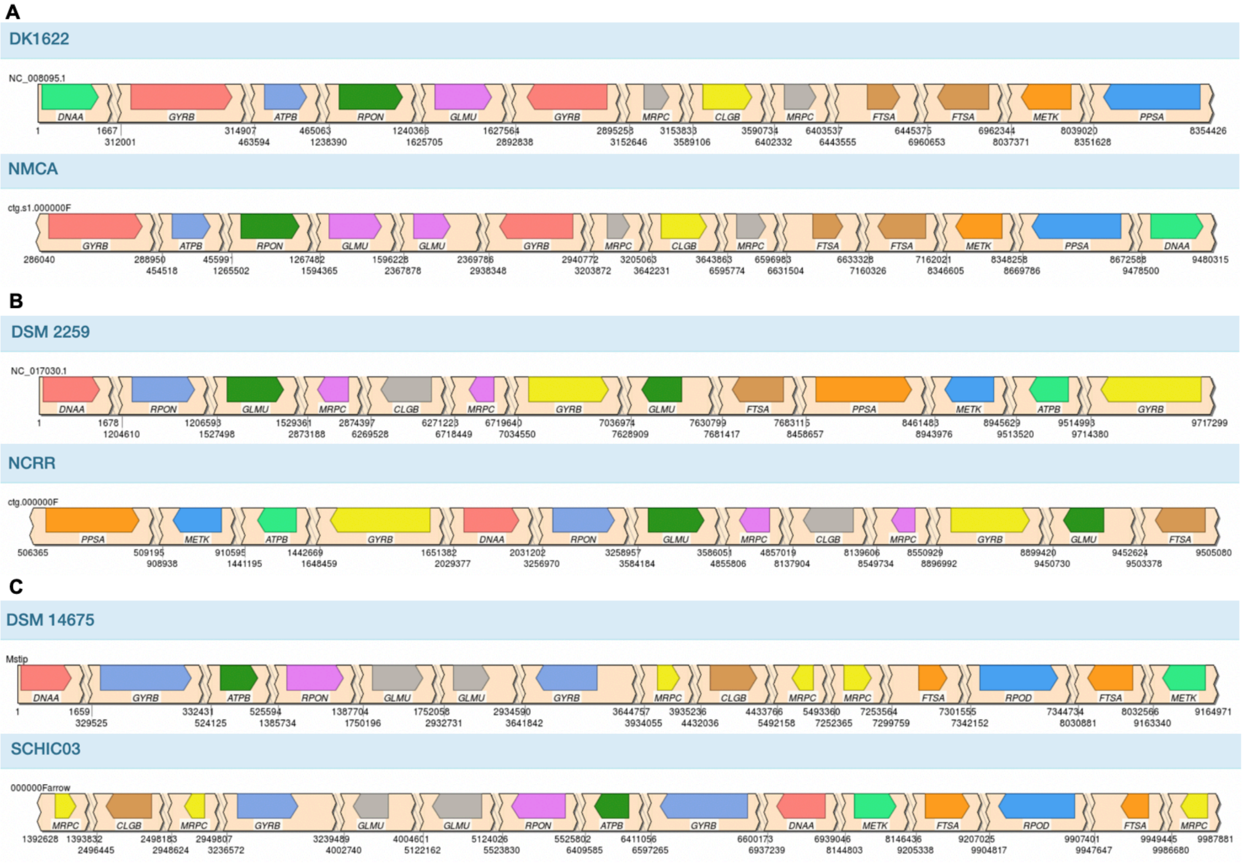


**Supplemental Figure 8:** Shared synteny of *atpB, clgB, dnaA*, *ftsA*, *glmU, gyrB, metK, mrpC, ppsA,* and *rpoN* between A) *M. xanthus* DK1622 and *M. xanthus* NMCA and B) *C. coralloides* DSM 2259$^{T}$ and *C. coralloides* NCRR. C) Shared synteny of *atpB, clgB, dnaA*, *ftsA*, *glmU, gyrB, metK, mrpC, rpoD,* and *rpoN* between *M. stipitatus* DSM 14675$^{T}$ and SCHIC03.

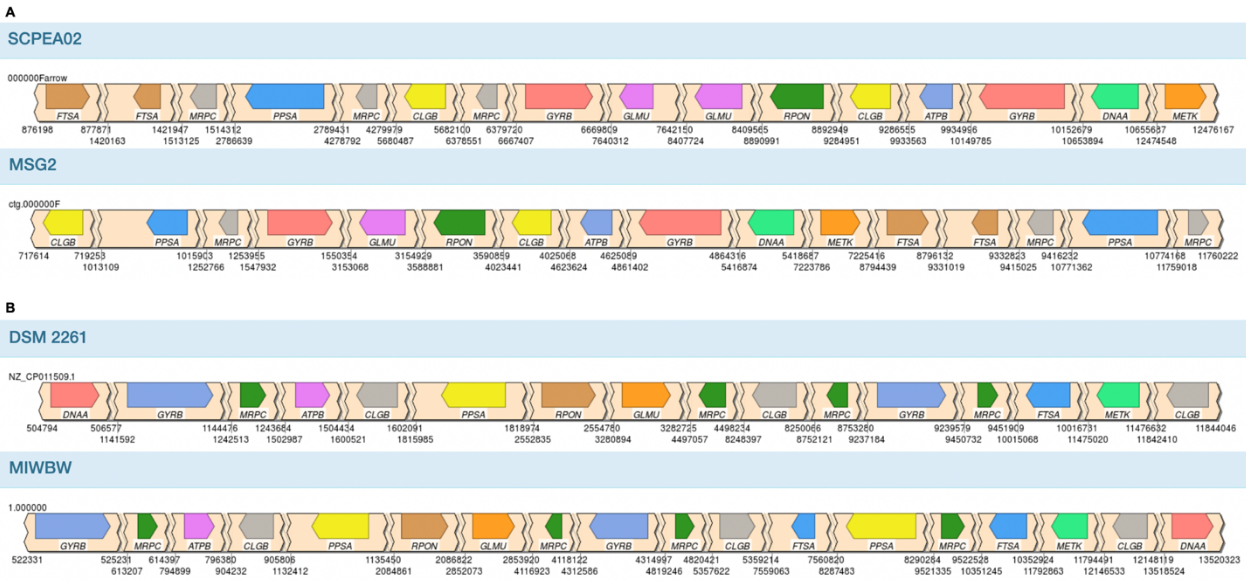


**Supplemental Figure 9:** Shared synteny of *atpB, clgB, dnaA*, *ftsA*, *glmU, gyrB, metK, mrpC, ppsA,* and *rpoN* between A) SCPEA02 and MSG2 and B) *A. gephyra* DSM 2261$^{T}$ and MIWBW.
